# Deep Learning-Meets-Active Surfaces: A Unified Framework for Multi-Object Segmentation and Shape Analysis in 3D Fluorescence Microscopy

**DOI:** 10.64898/2026.09.30.755627

**Authors:** Quentin Rapilly, Anaïs Badoual, You-Hsuan Liu, Charles Kervrann

## Abstract

In this paper, we introduce Nagini3D, a 3D biological image segmentation method that hybridizes two high-performing yet highly complementary segmentation paradigms: deep learning-based approaches and snake-based methods. Our approach leverages both the high efficiency of data-driven techniques and the convenient shape representation provided by active surfaces, which proves highly practical for incorporating a priori knowledge as well as for the post-analysis of segmented objects. By considering a complete 3D volume, our method ensures segmentation consistency across all three dimensions, unlike many approaches that handle volumes in a slice-by-slice manner. Additionally, it theoretically enables the representation of any object with spherical topology, overcoming the limitations of common 3D methods that rely on object convexity assumptions. Furthermore, the continuous representation of objects allows for a resolution-independent description, facilitating the extraction of various geometric features of interest (e.g., local curvature, principal deformations). We demonstrate both the quality of the segmentation produced by our method and the richness of the accessible geometric features through an extensive set of experiments conducted on multiple real and simulated datasets, including objects with challenging geometries.

---

The combined advancements in microscopy and digital technology have enabled the observation of cell mechanisms at increasingly smaller scales, as well as the storage and analysis of these now numerous observations. Indeed, the growing volume of data makes exhaustive analysis by specialists time-consuming and impractical. Furthermore, the acquisition of three-dimensional data complicates the task even further, as these data cannot be easily displayed on 2D computer screen.

To facilitate the analysis of these images, automated segmentation methods have emerged, first in 2D and later in 3D. Classical methods, based on active contours or morphological operations, were widely used for a long time but have now been almost systematically replaced by deep-learning-based methods due to their efficiency. However, active contours still sometimes offer significant advantages and provide capabilities that deep-learning approaches do not incorporate.

Throughout this paper, we present **Nagini3D**, a unified segmentation method that combines deep-learning with active surfaces. This approach merges the efficiency of deep-learning with the practicality of parametric surfaces for object representation, drastically simplifying *a posteriori* shape analysis.

## Classical methods

Methods for segmenting images have been around for decades, based on geometric descriptions of objects, morphological operators or statistical methods. For example, Yuille *et al*. [1] developed application-specific parametrized templates for shapes with well-defined a priori knowledge, such as facial features like eyes or lips. These models use simple 2D geometric representations. A complementary approach involves deformable templates, which model random deformations of an initial shape. Grenander *et al*. [2, 3] achieved early success in image segmentation by using statistical deformable models to capture local deformation patterns. Markov models and Monte-Carlo methods were later employed to estimate optimal deformations from image data [2, 3, 4, 5, 6]. Cootes et al. [7] introduced a data-driven method, learning both shape structure and deformation parameters from a set of representative examples, with applications in object tracking [5]. Staib and Duncan [8] further advanced the field by combining parametric snakes (B-splines) with Fourier decomposition for analyzing deformable biomedical structures. While these methods are robust to noise, their computational cost increases significantly when stochastic iterative optimization is required and no good initial estimate is available. Later, “*snakescules*” [9], integrated with fast algorithms and Markov point processes [10], have been developed for detecting cells or nuclei in fluorescence microscopy, building on these principles.

We rather focused on snakes, also called active contours in 2D, because of their wide use for microscopic images. This family of methods can be divided into three categories: (i) **points snakes**, assuming that the contour is described by an ordered set of pixel locations which are adjacent two by two [11, 12]; (ii) **geodesic snakes**, relying on the level sets of a continuous scalar function defined on the image domain [13, 14, 15]; (iii) **parametric snakes**, described by a curve parameterized by a small number of parameters, namely the “control points” [16, 17].

The latter has largely been used in 2D because parametric snakes are quite easy to optimize due to their small number of parameters and because they provide a continuous representation of segmented objects. Extensions using parametric surfaces instead of contours generalize this approach to 3D images [18, 19, 20]. Note that, the first approach that introduces fully parametric spline-based 3D snakes was only proposed in the past decade [21].

## Deep-learning methods

The advent of deep-learning methods in recent years has revolutionized the field of image analysis. Highly effective segmentation methods have emerged in conventional computer vision [22, 23], opening the gate for methods designed for biology and microscopy images. Concerning 2D microscopy images, there is no debate, Cellpose [24] (that predicts heat diffusion gradient used to regroup pixels belonging to the same object) and StarDist [25] (that localizes objects and represent them as star-convex polygons) outperform both the “classical”^1^ methods and their deep-learning competitors in terms of versatility, efficiency, and their widespread adoption within the biology community. Moreover, CNN/U-Net based methods [26] solved a major issue of snake algorithms that are not well suited for multi-object segmentation. Cellpose, has recently been updated to leverage the Segment-Anything-Model (SAM) [27] from Meta based on transformers, making Cellpose-SAM [28] more efficient than previous Cellpose models.

In 3D, no clear consensus has been established so far. Many 3D segmentation methods rely on 2D models applied to series of 2D slices. The resulting 2D segmentation masks are further merged to reconstruct a volumetric segmentation. Cellpose proposes its own merging algorithm averaging results obtained along each dimension. Other tools can be used to merge segmentation provided by any 2D methods such as uSegment3D [29] or Cell-Stitch [30]. However, some methods do process the entire 3D volume in a single pass. Despite the difficulty of labeling many 3D volumes, those methods focusing on the entire volume are in general more accurate when a dataset dedicated to the considered task is available. The lack of performance of methods based on 2D models often comes from the errors made by the aggregation algorithm while stitching the slices. The only purely 3D method used in practice by biologists is StarDist3D [31], the counterpart of StarDist [25], based on star-convex polyhedra. The alternative method Omnipose [32], that extends Cellpose approach to 3D, is used occasionally, but is not as popular as StarDist3D.

## Foundation models

Foundation models are increasingly prevalent in artificial intelligence, aiming to provide tools that deliver strong results across diverse scenarios without requiring task-specific training. However, foundation models dedicated to biological image segmentation are rare and recent. Cellpose-SAM [28] can be mentioned as one of the most famous as the extension of a largely used method (Cellpose [24]). It relies on the following features: i) its reliance on a Vision Transformer [33] (ViT); ii) the use of large training datasets (including pre-training on non-biological data). Other models have been regularly emerging (e.g., CellSAM [34], *µ*SAM [35]), but all remain focused on 2D images and rely on aggregation algorithms to process 3D images. Indeed, designing a fully 3D foundation model is barely feasible because: i) training a ViT on 3D images requires prohibitive computational resources; ii) annotated 3D data remains scarce. Consequently, 3D biological image segmentation typically relies on expert models dedicated to specific tasks.

## Contributions

Despite being often less accurate and primarily designed for single-object segmentation, active contour and surface approaches offer practical advantages: continuous contour description, easy *a posteriori* shape analysis, and prior shape knowledge. To our knowledge, previous deep-learning methods, which rely on pixel/voxel-based segmentation, do not support these features.

One naturally considers hybridizing both approaches. SplineDist [36] may be considered as a first approach to extend StarDist2D [25]. It uses spline-based parametric contours, thus providing continuous object representations while addressing a key limitation of StarDist: with contours, non-star-convex objects can also be represented. In [37], the authors compare different 2D shape models (spline, Fourier descriptors, etc) combined with CNNs. In [38], authors propose a variant of StarDist3D based on spherical harmonics as shape model, providing continuous object representations but it still handles star-convex shapes only.

In this paper we present **Nagini3D**, a fully 3D segmentation methods combining both deep-learning and active surfaces, to leverage the strengths of each. It can be considered as an extension of StarDist3D as SplineDist is in 2D microscopy, but includes additional feature outputs. The main contributions of the approach are the following ones:

- a paradigm to train a U-Net to both localize objects and predict a closed and smooth (*C*_1_) parametric surface representing each object. In particular, we introduce a loss function based on optimal transport and a regularization term on surfaces to guarantee plausible (non-self intersecting) shape predicted by the network;
- a two-stage segmentation method: the network output can either be used directly or enhanced via an optional optimization technique to minimize an user-defined energy functional, akin to conventional active surface models (snakes). The default energy available in the pipeline adapts to object density in a neighborhood and low/high contrast. It is well appropriate to accurately delineate boundaries of individual objects. As Nagini3D cleverly leverage GPU efficiency, it is the fastest method tested without optimization, and is comparable to competing methods with optimization;
- user-friendly functionalities for shape analysis of the detected objects, providing a direct access to mean local curvature or principal deformation of shapes.

All the code of the method is available on Github (https://github.com/QuentinRapilly/NAGINI-3D/) with tutorials for training and testing. A Napari plugin has been developed to make the method as user friendly as possible for biologists. It can be installed with one single line of code and is designed to run on any computer but is fasten using a GPU. The plugin description is available at https://github.com/QuentinRapilly/nagini3d-napari.

## Results

### Overview of Nagini3D

**Nagini3D** aims to segment multiple objects in microscopy volumes with high accuracy and robustness. To achieve this, it combines the computational efficiency of deep-learning with parametric surface object representation. Given an input image volume, a 3D U-Net architecture predicts two types of channel maps: i) a score map, used to localize objects of interest within the volume; ii) surface parameter maps. For any given spatial location, the values extracted from the surface parameter maps enable, the generation of a parametric surface delineating the object to which the location belongs to. Consequently, the final segmentation is achieved by generating the predicted surfaces at each estimated object position. The training procedure consists in comparing the predicted score map to a ground-truth (GT) one and to match the predicted surfaces with object samplings both derived from a GT mask. The pipeline is illustrated in Fig. 1 and the technical details are provided in the Method Section.

**Figure 1.**
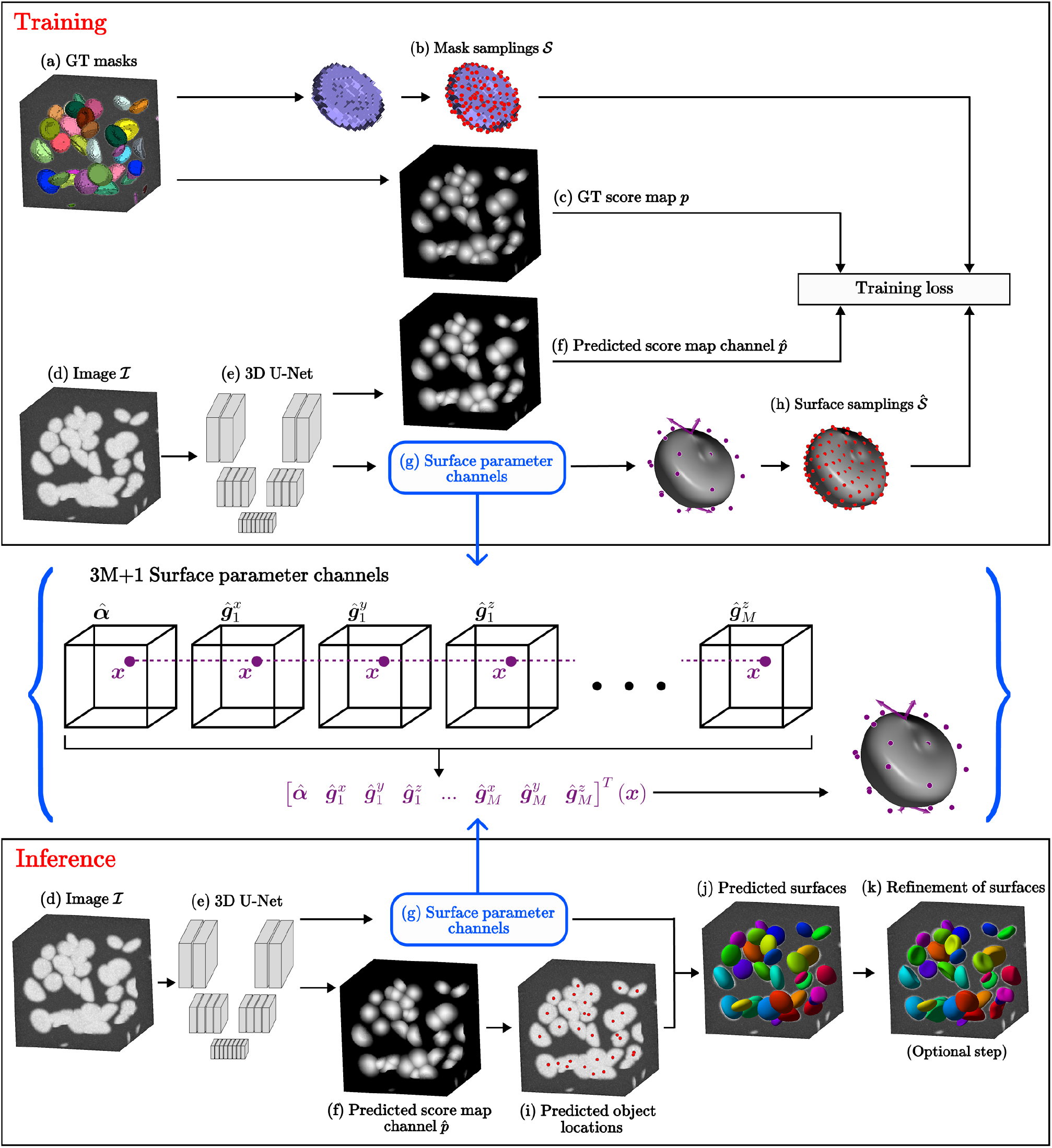
Overview of Nagini3D. During the model **training**, the ground-truth (GT) mask **(a)** allows to generate a GT sampling *S* for each object contained in the patch **(b)** and GT score-map ***p* (c)**. The image patch *I* **(d)** is given to the network **(e)** that predicts i) a score map 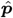 **(f)**; ii) surface parameter channels **(g)**. For any given position ***x*** in the patch, each channel provides one of the parameter coordinates generating a surface that represents the object to which ***x*** belongs to. The predicted surfaces are sampled **(h)** and matched with the GT samplings. The GT and predited score maps are compared. See equation (4) for details. During the **inference:** the score map and the parameter coordinates are predicted identically than for training. The score map 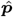 allows to localize objects **(i)** and the surface representing the object are generated at this location **(j)**. An optional snake-based refinement can be applied to enhance the result **(k)**.

The key advantage of **Nagini3D** is that the continuous representation of objects allows a straightforward geometric analysis of segmented objects (PCA, curvature, …). The method achieves state-of-the-art performance and is particularly effective on challenging shapes. It generates accurate results with a few annotated volumes only. The shape model used permits to store the segmentation in a memory-efficient manner and facilitate the simulation of phantom datasets.

### Nagini3D allows straightforward geometric analysis of segmented objects

The major advantage of **Nagini3D**, compared to current state-of-the-art approaches in 3D biological image segmentation, is its ability to provide a continuous representation of each segmented object (see Fig. 2(b-c)), rather than just a discrete voxel-based mask limited by the resolution of the input image. This feature, achieved through the parametric surface representation of objects (see Fig. 2(a)), significantly simplifies post-segmentation analysis.

**Figure 2.**
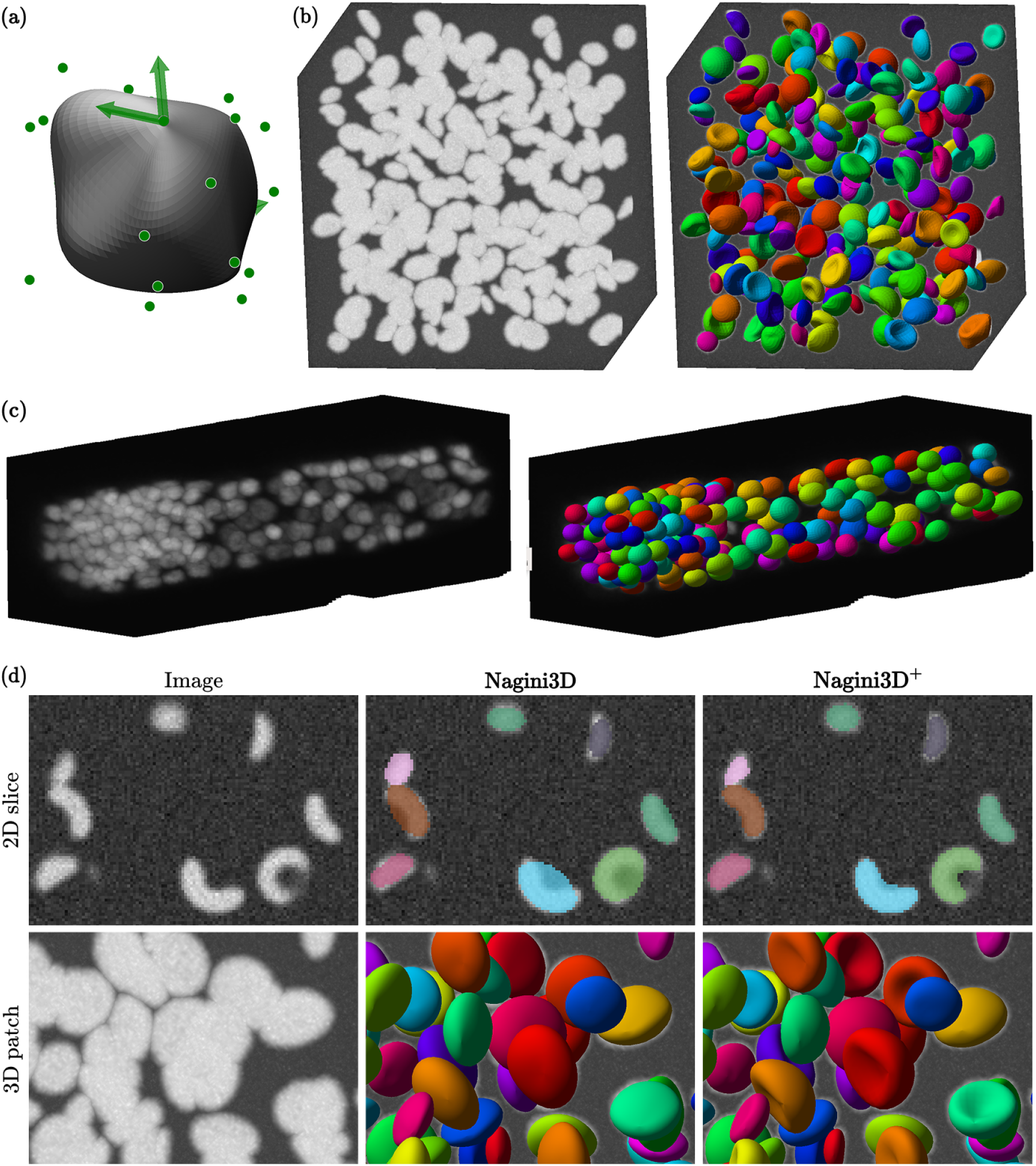
Nagini3D predicts continuous object representations. **(a)** Surface parameterization used in **Nagini3D**: green dots are the control points and green arrows are the normal at poles. **(b-c)** 3D volumes and the corresponding surfaces predicted by **Nagini3D**. Volume **(b)** is synthetic and represent non-star-convex objects (CAPS dataset). Volume **(c)** represent a *C*.*Elegans* larva imaged using fluorescence microscopy. **(d)** Influence of the refinement process on surfaces predicted on a volume of CAPS (see Section “Nagini3D precision is enhanced by object-wise optimization” for details on refinement step). The segmentation are overlaid on the raw images.

Actually, **Nagini3D** provides a direct access to objects scale, and for the first time multiple shape features such as mean or Gaussian local curvature. In Fig. 3(a), the mean local curvature is displayed on objects segmented in a volume from CAPS dataset^2^. It enables a straightforward way to cluster and classify object based on morphological characteristics (e.g., presence of concave areas).

**Figure 3.**
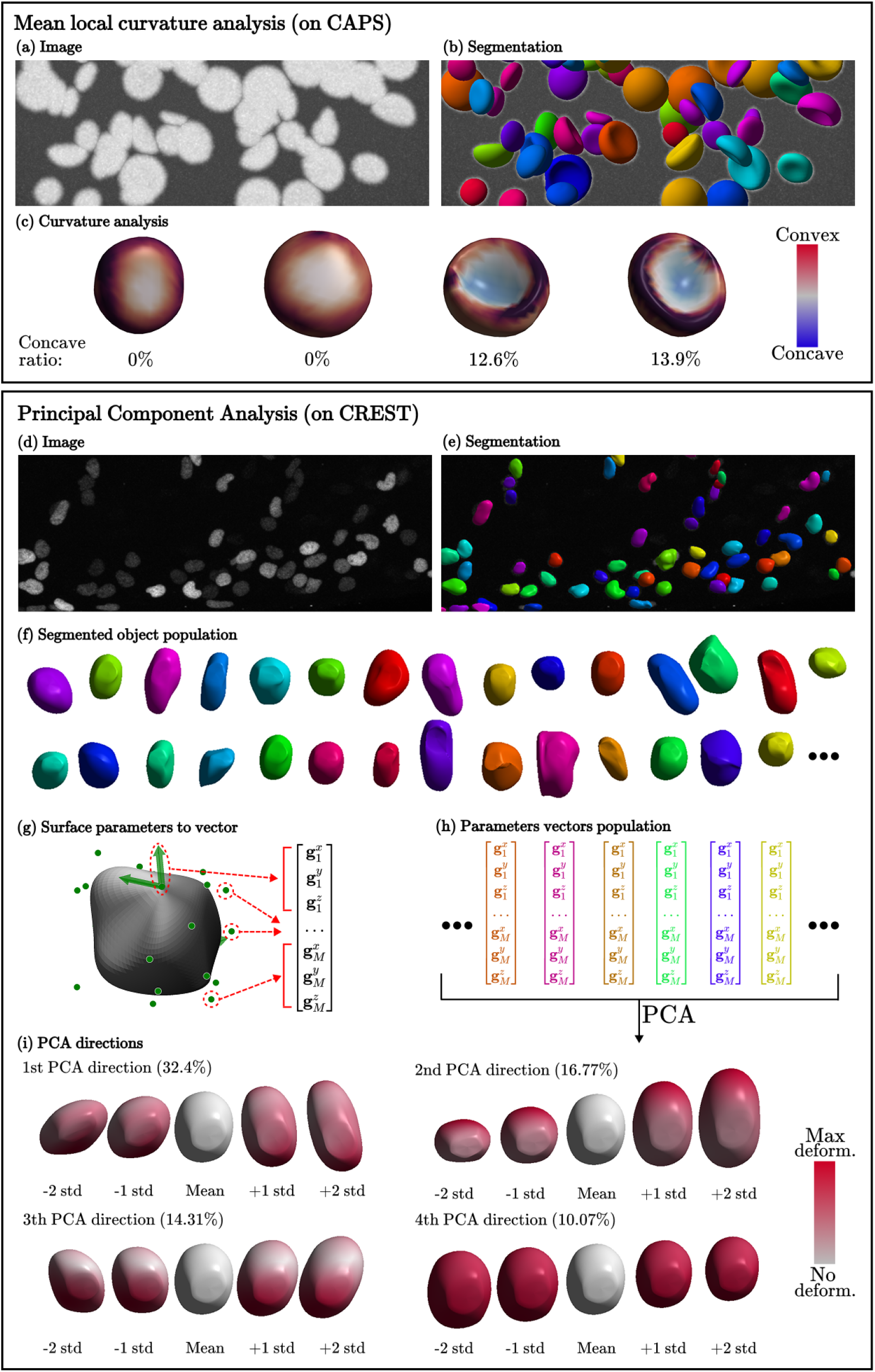
Analysis of shape variability: local curvature and statistical deformation modes. **Local curvature:** a volume from CAPS set **(a)** is segmented **(b)**. The mean local curvature can computed on predicted objects **(c)**. The convex parts are in red, the concave ones in blue. The proportion of concave areas of the surface allows to cluster shapes (left: no concavity; right: important concavity). **Principal component analysis:** a volume from CREST **(d)** is segmented **(e)**. For each object in the object population **(f)**, one can generate a vector with the surface parameter coordinates **(g-h)**. A PCA is performed on these vectors. The four principal directions are displayed **(i)**.

Principal Component Analysis (PCA) can also be performed directly on surface parameters (3D coordinates of the control points, poles and normal at poles) to obtain valuable insights into deformation patterns across the segmented population in a given volume or set of volumes. In Fig. 3(b), PCA was performed on the surface parameters of objects extracted from the CREST set (zebrafish neural crest cells). The first principal directions reveal that the dominant deformation is cell elongation as expected. In this example, cells migrate through narrow channels, they are constrained to elongate along that direction^2^.

Using **Nagini3D**, these valuable shape analysis features can be computed directly and instantly after segmentation, whereas traditional segmentation methods typically require sophisticated time-consuming intermediate steps and use of external heterogeneous software.

### Nagini3D precision is enhanced by object-wise optimization

To refine the network prediction and ensure that the predicted surfaces accurately delineates object contours, an individual surface optimization step can be applied. Specifically, as our network outputs surface parameters that are rigorously the same than those used in conventional active surfaces algorithm, **Nagini3D** enables (optionally) to optimize each predicted surface by the network by minimizing an energy functional based on image features (e.g., intensity gradient, etc.), akin to conventional snake algorithms. This two-step approach then combines a global and an individual strategy. First, the network is trained to predict all surfaces simultaneously; and second, each surface is independently refined based on image features, ensuring the highest possible accuracy and correcting potential errors introduced by the network simultaneous prediction of multiple surfaces. As individual adaptation may be less desirable or even counterproductive in dense regions since it can deteriorates reliable predictions due to the poorly defined boundaries of closely adjacent or weakly contrasted objects, **Nagini3D** then automatically adapts the level of refinement depending on image contrast and object density in the surface vicinity. Details on the energy chosen for this step are provided in the Method section.

In what follows, the prediction output from the network is referred to as **Nagini3D**, while the refined prediction obtained through this additional optimization step (which can be used optionnaly) is denoted as **Nagini3D**^+^.

Figure 2(d) illustrates the effect of the refinement step on the predicted segmentation. The impact on the masks in terms of local details is noticeable, especially for complex shapes: the initial segmentation is coarse and does not segment accurately the concavity of the objects in CAPS set. After refinement, the concavities are correctly segmented, which explains the important improvement in terms of Boundary Intersection-over-Union (BIoU) in Table 1.

**Table 1.** Detection and segmentation performance measures of tested methods. The recall and precision are both computed with *TP* (0), *FP* (0) and *FN* (0) (i.e., *τ* = 0) to assess the quality of the detection. The score is the integral of the *acc*(*τ* ) for *τ ∈* [0, 1]. The mean/med IoU and BIoU are given among the TP to assess the quality of the segmentation for the objects correctly detected. Higher score is better for all metrics except for inference time.

|  | Nagini3D | Nagini3D<sup>+</sup> | SD3D | OP3D | CP-SAM | CP2.5D | uSeg2.5D | StitchCP2D | StitchSD2D |
| --- | --- | --- | --- | --- | --- | --- | --- | --- | --- |
| **BONES** |  |  |  |  |  |  |  |  |  |
| Recall | 0.951 | 0.949 | 0.965 | 0.984 | **0.987** | 0.955 | 0.888 | 0.930 | 0.976 |
| Precision | **0.899** | **0.899** | 0.882 | 0.356 | 0.764 | 0.809 | 0.325 | 0.755 | 0.593 |
| Score | 0.565 | 0.659 | **0.669** | 0.253 | 0.500 | 0.442 | 0.099 | 0.346 | 0.318 |
| Mean IoU | 0.673 | 0.791 | **0.793** | 0.717 | 0.680 | 0.584 | 0.313 | 0.511 | 0.555 |
| Med. IoU | 0.697 | **0.849** | 0.828 | 0.780 | 0.722 | 0.595 | 0.318 | 0.545 | 0.576 |
| Mean BIoU | 0.535 | **0.725** | 0.703 | 0.615 | 0.578 | 0.428 | 0.130 | 0.380 | 0.463 |
| Med. BIoU | 0.555 | **0.782** | 0.733 | 0.654 | 0.613 | 0.417 | 0.114 | 0.387 | 0.476 |
| **CAPS** |  |  |  |  |  |  |  |  |  |
| Recall | **1.000** | 0.998 | **1.000** | **1.000** | 0.996 | 0.991 | 0.974 | 0.979 | 0.983 |
| Precision | **1.000** | **1.000** | 0.958 | 0.162 | 0.977 | 0.976 | 0.996 | 0.905 | 0.729 |
| Score | 0.753 | **0.865** | 0.730 | 0.114 | 0.772 | 0.754 | 0.578 | 0.670 | 0.487 |
| Mean IoU | 0.752 | **0.859** | 0.779 | 0.688 | 0.788 | 0.773 | 0.592 | 0.764 | 0.692 |
| Med. IoU | 0.752 | **0.868** | 0.816 | 0.730 | 0.798 | 0.780 | 0.594 | 0.794 | 0.741 |
| Mean BIoU | 0.600 | **0.763** | 0.696 | 0.589 | 0.654 | 0.627 | 0.316 | 0.643 | 0.583 |
| Med. BIoU | 0.582 | **0.770** | 0.722 | 0.618 | 0.660 | 0.632 | 0.314 | 0.666 | 0.622 |
| Mean inf. time (s) | **4.75** | 11.76 | 8.35 | 7.86 | 11.27 | 27.20 | 34.31 | 62.96 | 59.72 |

### Nagini3D is fast and achieves state-of-the-art competitiveness

**Nagini3D** was benchmarked against state-of-the-art methods across real and simulated 3D datasets of fluorescence microscopy images. Complete details on evaluated algorithms (training and/or inference settings), datasets (train/val/test split, images size, number of object per image) and metrics are provided in the Method section. The average inference time and the detection and segmentation scores are summarized in Table 1. The accuracy curves of the evaluated methods are displayed in Fig. 4(a).

**Figure 4.**
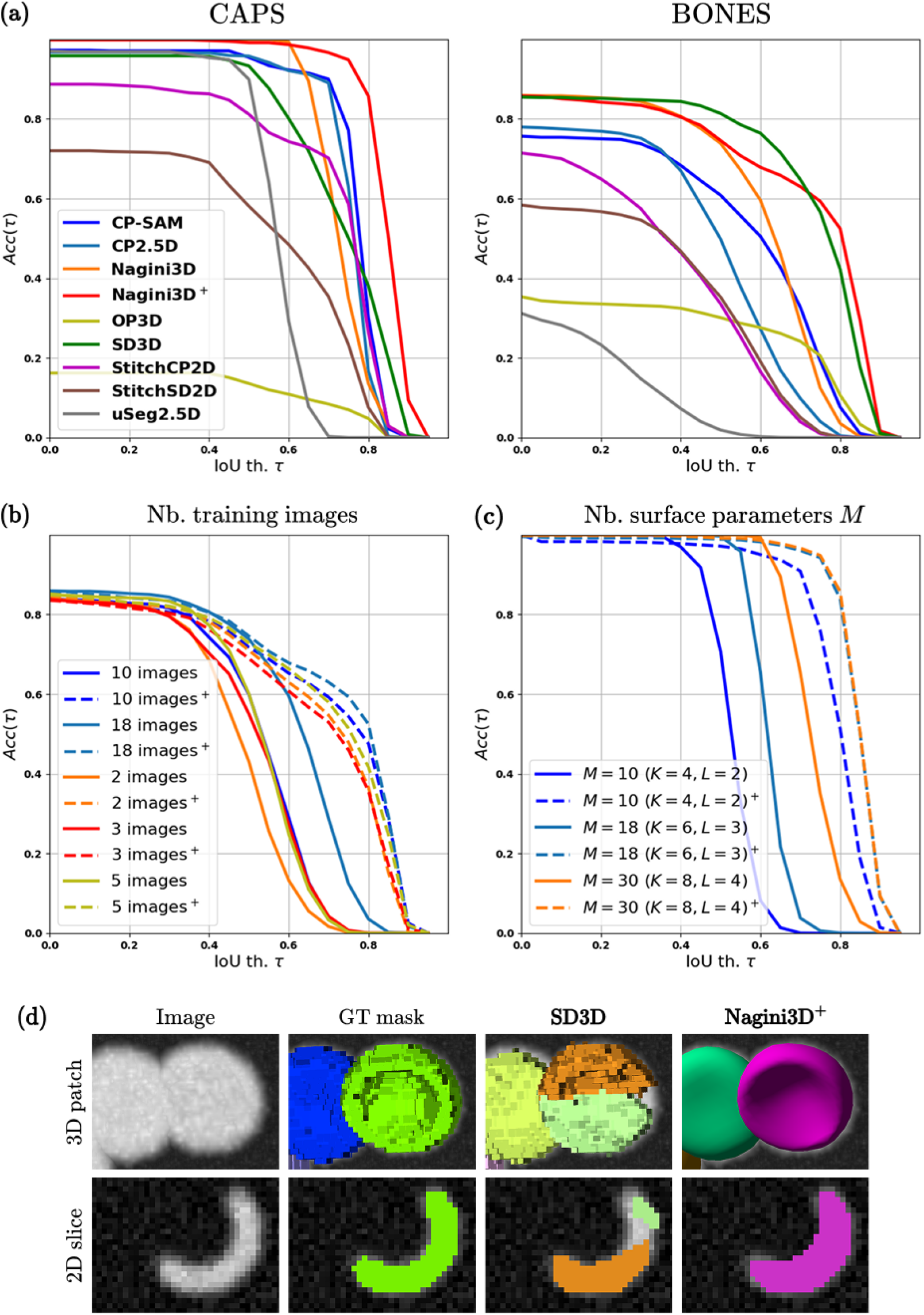
Performance comparison with accuracy curves. **(a)** for each evaluated methods over CAPS (left), a set of synthetic images depicting non-star-convex objects; and BONES (right), a real fluorescence microscopy image set. The evolution of **Nagini3D** and **Nagini3D**^+^ performance depending on the amount of training images used (BONES set) is illustrated on **(b)**; while **(c)** represents the influence of the number of surface parameters (CAPS set). For **(b)** and **(c)**, dashed lines correspond to refined results (**Nagini3D**^+^) corresponding to plain lines (**Nagini3D**). **(d) The non-star shape case:** difference of behavior for non-star-convex objects. **SD3D** cannot represent such objects and split them into two parts. **Nagini3D**^+^ is not limited by convexity concerns.

On BONES, the 3D methods outperform the 2.5D algorithms in terms of both detection and segmentation accuracy. The exception is **OP3D** [32], which, despite producing highly accurate masks, tends to generate false positives (FP) due to residual pixel groups or splitting nuclei into multiple labels, as reflected in its low precision. The best performing methods are **SD3D** [31] and **Nagini3D**^+^ with very similar performance in detection and segmentation. The ranking of the best methods varies slightly depending on whether the IoU or BIoU (more sensitive to boundary errors) is considered. **Nagini3D** appears to segment object boundaries more accurately as the performance gap between **SD3D** [31] and **Nagini3D**^+^ slightly increases using the BIoU instead of IoU.

On CAPS, the difference between 3D and 2.5D algorithms is less noticeable: **CP2.5D** [24] and **CP-SAM** [28] achieve detection and segmentation scores comparable to 3D methods. Unlike **SD3D** [31], whose performance drops to that of the best aggregation algorithms, **Nagini3D**^+^ still outperforms all other methods in terms of both detection and segmentation. This difference can be explained by the presence of non-star-convex objects in this dataset: **SD3D** [31] cannot represents such objects and split them into several parts while our approach is not limited by such shapes. This difference of behavior is longer discussed in Section “Nagini3D better handles objects with complex shapes”.

In addition, according to last row of Table 1 (reporting the mean inference time per method on CAPS test set), **Nagini3D** is the fastest among all the evaluated methods, and the inference time is of the same order even when the refinement step is applied (**Nagini3D**^+^). This can be explained by our fast local maxima extraction based on convolution and details in Supplementary Section.

### Nagini3D better handles objects with complex shapes

As mentioned in Section “Nagini3D is fast and achieves state-of-the-art competitiveness”, **Nagini3D** outperforms its competitors when dealing with objects whose shapes deviate significantly from simple ovoids and whose convexity substantially differs from star-convex objects. Its direct competitor in the fully 3D methods category, **SD3D** [31], relies on star-convex polyhedron, an object representation that is intrinsically incompatible with non-star-shaped objects. **Nagini3D** has been especially designed to address this issue: the spline-based parametric surface used to represent object is convexity-agnostic and can thus represent any object homeomorphic to a sphere. As illustrated on Fig. 4(d), when **SD3D** [31] is used to segment non-star-convex objects, these are split in multiple parts, leading to false-positive predictions and under-segmented objects. On the contrary, **Nagini3D**^+^, represent those objects using a single surface and therefore get almost perfect detection scores and the best mean IoU by a noticeable margin on the the **CAPS** set containing non star-convex objects.

To convince the reader of **Nagini3D**^+^ effectiveness in segmenting objects with challenging geometries, Fig. 5 displays visual segmentation results from the three top-performing methods (**CP-SAM, SD3D**, and **Nagini3D**^+^ as identified in Section “Nagini3D is fast and achieves state-of-the-art competitiveness”) on CREST, a dataset featuring zebrafish trunk neural crest cells migrating through confined environments creating nuclear deformation. The objects are sometimes deformed to the point of approaching non-star-convex shapes (see Fig. 5) and shape analysis is crucial for biologist interested in those cells.

**Figure 5.**
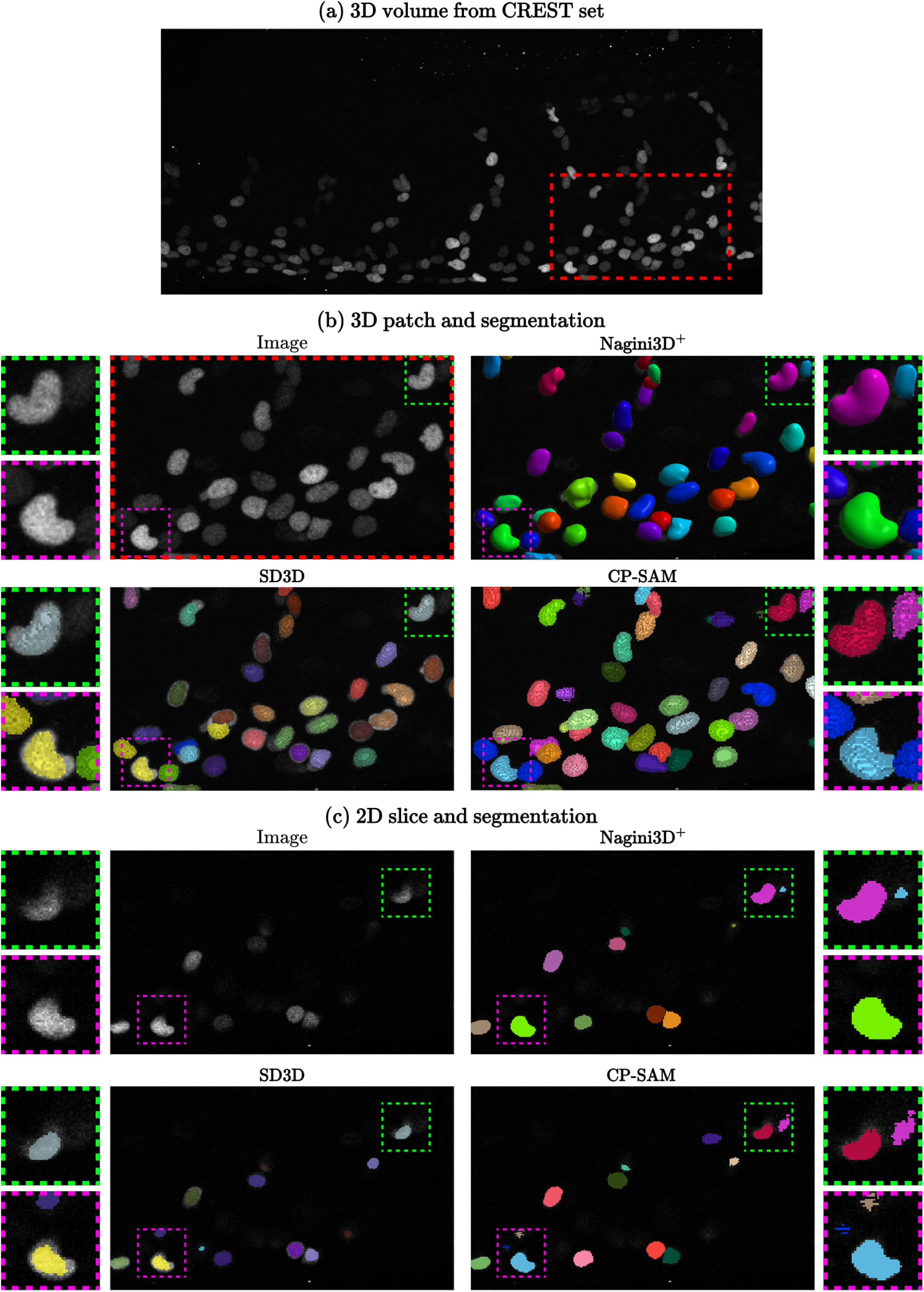
Performance of methods on variable (non-convex, and non-star convex) shapes. Example of volume from CREST set **(a)**. The volume contain challenging 3D “kidney-shaped” objects. The segmentation obtained using **Nagini3D**^+^, **SD3D** and **CP-SAM** are displayed for a small 3D patch **(b)** and for a 2D slice **(c)**. While **Nagini3D**^+^ consistently segments the entire object, **SD3D** tends to under-segment (particularly challenging objects) and **CP-SAM** does create undesired masks and sometime generates discontinuous segmentation masks.

On that challenging set, **Nagini3D**^+^ segments correctly the majority of the objects, while **SD3D** under-segment multiples, particularly the kidney-shaped objects. That underline another positive aspect of **Nagini3D**^+^: for that set, the GT masks used for training may not perfectly align with object boundaries, sometimes being shifted a few voxels inward. While **SD3D** mimic the defaults of the GT masks, **Nagini3D**^+^ adapts thanks to the refinement step, that corrects such errors. **CP-SAM** segmentation masks are also of great quality but some are not continuously merged and a few small false-positive fragments occur. These results show that **Nagini3D**^+^ is of particular interest if the GT masks are not perfectly aligned with the object to segment and when challenging non-star-convex shaped objects are present in the set.

### Nagini3D only needs few annotated samples

Exhaustive annotation of multiple 3D volumes (every object labeled) is time-consuming. While **Nagini3D** requires such labels for training^3^, it offers two key advantages due to its two-step inference process. First, the method performs efficiently even with a few annotated volumes used for training: Fig. 4(b) shows accuracy curves for a model with identical parameters trained on increasingly smaller subsets of the BONES dataset. The results demonstrate that segmentation quality improves with the amount of annotated data used during training, particularly for post-network results. However, after refinement, the model trained on just two images (corresponding to approximately 450 labeled objects) already performs very well and is close to the model trained on the full dataset (+4000 labeled objects).

Additionally, the data can be annotated coarsely rather than finely (with imprecise boundaries), for example, using a fast automatic method that requires no training. As mentioned in Subsection “Nagini3D better handles objects with complex shapes”, the refinement step corrects potential errors inherited from training. Thus, Nagini3D is an excellent tool for using a fully 3D model while minimizing annotation requirements, saving time and achieving highly satisfactory results with minimal effort.

A detailed tutorial on annotation requirements is provided in Supplementary Section .

### Nagini3D leverages a memory-efficient representation of objects

Using **Nagini3D**, object in an image can be represented with only a few parameters, making it a memory efficient data representation. Storing 3D data can be quite heavy and for large image, a segmentation mask can represent several hundred of mega-bytes (MB). A convenient way to save storage is to save the location and the shape parameters only and generate segmentation masks on the fly if required. Besides, for all of the shape analysis features described in Section “Nagini3D allows straight-forward geometric analysis of segmented objects”, generating the masks is not even required: surface and shape analysis can be displayed using Napari with the shape parameters only. As an example, a segmentation mask for larger images of BONES set weights 90MB^4^ while storing the surface locations and parameters weights 0.2MB. This quantity will slightly vary depending on the amount of surface parameters used but the representation capability is good even with a limited number of shape parameters as described in Fig. 4(c). With *M* = 18 surface parameters

(leading to 54 = 3 *×* 18 parameter coordinates), the model reach its performance threshold and the quality of **Nagini3D**^+^ no longer increase if *M* is increased to 30. For very sophisticated shapes, a larger amount of parameters could be required, but a trade-off with the computation capability required to train the model should be found.

Moreover, the set of collected object surface parameters can be used to constitute a dictionary for phantom benchmark simulations, which is a more convenient way than conventional voxelwise maps produced by by usual deep-learning methods.

## Discussion

**Nagini3D** is a competitive segmentation method that achieves state-of-the-art performance and simplifies shape analysis, making it both convenient and straightforward. It can be used for local-curvature analysis and conventional PCA-based analysis; paving the way for more sophisticated analysis (e.g., including optimal transport) could be explored in future work. These proposed shape analysis functionalities are the pillars of mechanobiology, which studies the impact of physical forces on cells and tissues. The surface representation could also be used to facilitate tracking and estimate object deformation over time, similar to the analysis performed in [39, 40, 41] for single object tracking. The possibility of incorporating shape priors (either during refinement or in the training loss to facilitate training) has not been explored so far but could be highly beneficial. Finally, the Napari plugin, which makes **Nagini3D** user-friendly for end users, will be regularly updated with new features (e.g., enabling users to manually adjust surfaces by moving control points). The object representation makes the storage of the segmentation memory-efficient and segmented objects can be used afterwards for data generation as a wide range of geometrical transformations can be applied to make data-augmentation.

One key limitation of our method is that the shape model struggles with very elongated objects and would require a large number of control points to accurately capture the shape. A potential solution is to use an alternative surface model (e.g., tubular parametric surfaces). Moreover, training our model required significant computational resources, as each patch is treated as a 3D volume. Depending on the dataset, GPU training memory requirements can range from 40 to 80 GB^5^, requiring a trade-off between batch and patch size and the number of surface parameters used. Fortunately, inference requires significantly less memory and can be performed on a standard laptop.

## Method

**Object representation** In **Nagini3D**, objects are represented with parametric spline surfaces with a spherical topology (see [21]) described as:

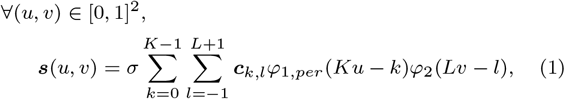

where *σ ∈* ℝ_+_ is an object scale factor, *{****c***_*k,l*_*}*_0*≤k≤K−*1;*−*1*≤l≤L*+1_ *∈* (ℝ^3^) ^Z*×*Z^are the 3D control points of the shape. The parameters *K* and *L* determine the number of control points and therefore the complexity of the surface. The basis functions *φ*_1,*per*_ and *φ*_2_ enforce various properties to the surface (smoothness, reproduction properties, …). The functions *φ*_1,*per*_ = *ϕ*_*K,per*_ and *φ*_2_ = *ϕ*_2*L*_ enable to reproduce ellipsoids. The function 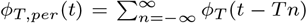 is the *T* periodic version of an exponential B-Spline *ϕ*_*T*_ (see Supplementary).

An alternative set of parameters, derived from the control points describes any closed and smooth shape with a spherical topology by satisfying closure and smoothness constraints (see Supplementary ). This set is:

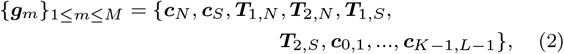

where ***c***_*N*_, ***c***_*S*_ denote the north and south poles of the surface respectively, and the vectors ***T***_1,*N*_, ***T***_2,*N*_ *∈* ℝ^3^, (***T***_1,*S*_, ***T***_2,*S*_ *∈* ℝ^3^, respectively) characterize the tangent plane to the north pole (south, respectively). The set contains *M* = *K*(*L−* 1) + 6 parameters including a subpart of the control points *{****c***_*k,l*_*}* _0*≤k≤K−*1;1*≤l≤L−*1_. For a surface with a spherical topology, the variables *u* and *v* represent the longitude and the latitude parameters of the surface. Equation (1) is finally rewritten using the *{****g***_*m*_*}*_1*≤m≤M*_ set as follows:

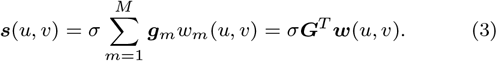

In Fig. 2(a), an example of surface parameterized using Equation 3 is represented.

Complete details on equations linking: i) *{****g***_*m*_*}*_1*≤m≤M*_ to *{****c***_*k,l*_*}*_0*≤k≤K−*1;1*≤l≤L−*1_; ii) basis functions *φ*_1,*per*_ and *φ*_2_ to (*u, v*) *→ w*_*m*_(*u, v*), *∀ m ∈ {*1, …, *M}* can be found in Supplementary .

**Network architecture** In **Nagini3D**, a CNN is trained to predict, in a single forward pass, an optimal set of parameters 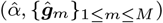 for each object in an image describing its boundaries. The CNN architecture we consider is a U-Net [26].

The input of the network is a 3D image ***I*** of size (*C*_*in*_, *H, W, D*), where *H, W* and *D* denote the height, width and depth of the image, respectively, and *C*_*in*_ is the number of channels (*C*_*in*_ = 1 in our experiments). The network predicts a volume of size (*C*_*out*_, *H, W, D*). For each voxel of the input image ***x*** = (*x, y, z*) *∈* Ω where Ω = *{*0, …, *H −* 1*} × {*0, …, *W −* 1*} × {*0, …, *D −* 1*}* is the voxel space, the network outputs *C*_*out*_ = 3*M* + 2 channels.

The first channel corresponds to a score map 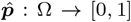 whose local maxima are assumed to represent object positions in the volume. It is used for object detection stage during inference. The 3*M* + 1 remaining channels are surface parameters maps: i) the object scale factor map 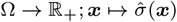; ii) the maps providing the three spherical coordinate of the *M* parameters Ω *→* (ℝ^3^)^*M*^ ; ***x*** ↦ *{****ĝ*** _*m*_(***x***)*}*_1*≤m≤M*_ of a surface centered at voxel ***x*** and delineating the object to which ***x*** *∈* Ω belongs; the vectorized representation of the surface parameter coordinates is 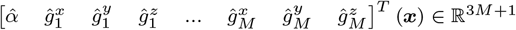.

### Training loss

The loss computed over a patch to train the network is defined as the average of a per-voxel term given for each voxel ***x*** *∈* Ω by:

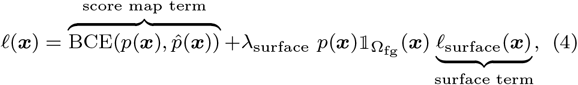

where BCE stands for Binary Cross Entropy. Here, *p*(***x***) and 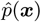 are the GT and predicted score at voxel ***x***, and respectively, 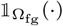 denotes the indicator function of the foreground voxels. The parameter *λ*_surface_ weights the importance given to the surface loss *ℓ*_surface_. This term is computed only on the foreground voxels (i.e., ***x*** *∈* Ω_fg_), is weighted by the GT score *p*(***x***) (to give more importance to the voxels that are more likely to be voxels of interest), and is defined as follows:

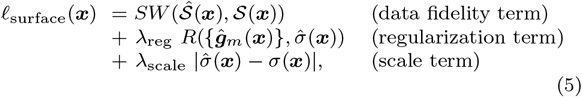

where *λ*_reg_ and *λ*_scale_ weight respectively the regularization term *R*(*·*) and the scale term. The surface loss compares the points 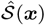 sampled on the surface predicted at ***x*** with a GT sampling *S*(***x***), using the Sliced Wasserstein Distance (*SW*). The scale term compare Details on each terms are given in following paragraphs. The GT scale is defined as 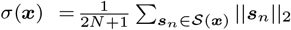, where ***s***_*n*_ *∈* ℝ^3^, (1 *≤ n ≤* 2*N* + 1) is a sampling point belonging to *S*(***x***). In all the experiments, the parameters are set to: *λ*_surface_ = 1, *λ*_reg_ = 0.01, *λ*_scale_ = 0.2.

### Score map term

The GT score map ***p*** is generated as follows: we set *p*(***x***) = 0 if ***x*** is a background voxel (***x*** *∈* Ω_bg_). If ***x*** is inside an object of index *j* (***x*** *∈* Ω), 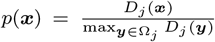 where 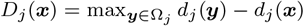 and *d*_*j*_ (***x***) = ||***b***_*j*_ *−* ***x***||_2_ is the Euclidean distance to the barycenter ***b***_*j*_ of the object Ω_*j*_ . Thus, *p*(***x***) is positive, increases as ***x*** gets closer to the barycenter of an object of interest, and reaches a maximum value of 1 if *∃j*, ***x*** = ***b***_*j*_ . For particular shapes, when the barycenter does not belong to the interior of the object, it is maximum for the point in this object that is the closest to the barycenter. Examples of GT score maps are shown in Supplementary Fig. 8(b).

### Surface fidelity term

To compare a predicted surface and a binary mask, both are sampled and the resulting point clouds are compared, following the principle described in [42] in 2D.

#### Predicted surface sampling

An even sampling of 2*N* + 1 points 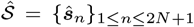, of a predicted surface is obtained by evaluating Equation (3) at longitudes and latitudes 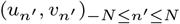 given by a Fibonacci lattice [43] (details in Supplementary ):

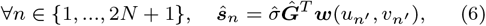

where *n*^*′*^ = *n − N −* 1. Once values of *N* and *M* are chosen (i.e., *K* and *L* fixed), vectors 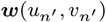, *∀ n*^*′*^ *∈ {−N*, …, *N}*, which are identical for all surfaces can be pre-computed and stored into a look-up table, leading to an efficient way to sample any surface by computing a weighted sum of the parameters. The simultaneous computation of multiple samplings become very fast, as it can be parallelized on GPU architectures.

#### GT contour sampling

The 3D boundary of a segmentation mask is extracted using morphological operation: the voxels lying in the difference between the original mask and its dilated version correspond to the boundary. Then, a “Farthest Point Sampling” (FPS) algorithm [44] is applied to uniformly sample the contour. The process begins by randomly selecting an initial point on the contour, followed by iteratively adding the farthest remaining point from the already sampled set until the desired number of points is achieved. It leads to a set of 2*N* +1 points *S* = *{****s***_*n*_*}*_1*≤n≤*2*N*+1_ evenly distributed on the GT contour. Each GT object is sampled once and the result is stored into a look-up table. Examples of samplings generated using FPS are shown in Supplementary Fig. 8(c).

#### Samplings matching

The Sliced Wasserstein Distance (SW) [45] is used to compare the predicted and GT samplings. This computationally efficient variant of the Wasserstein Distance (WD) can be defined as follows:

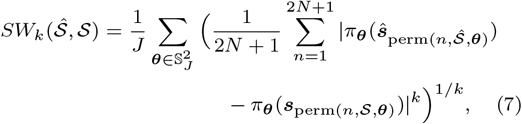

where 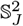 contains *J* random elements, uniformly sampled in the unit sphere S^2^. Here, *π*_***θ***_(***x***) = ***θ***^*T*^ ***x*** is the projection of ***x*** on the 1D subspace of ℝ^3^ directed by ***θ*** and the permutation perm(*·,χ, θ*) sorts the set of projections of elements of *χ* along direction ***θ***, *{π*_***θ***_(***x***_*n*_), ***x***_*n*_ *∈ X}*, in increasing order.

According to [46], formulation from Equation 7 correctly approximates the usual integral over S^2^ of the SW when *J ≥* 100. In Nagini3D, *J* is set to 100 and *k* to 1 (i.e., *SW* corresponds to *SW* _1_).

### Surface regularization term

Surface matching based on the SW does not penalize biologically unrealistic shapes (e.g., self-intersecting surfaces). To prevent them, the loss includes a regularization term *R*(*·*) penalizing the norm of surface derivatives as illustrated in Supplementary Fig. 10. This term is defined as:

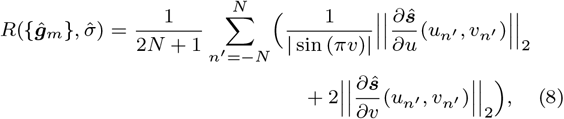

where 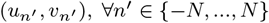 are the 2*N* +1 latitudes and longitudes given by the Fibonacci lattice. The surface derivatives are normalized to take into account the non-constant arc length of the surface parameterization. The surface derivative with respect to *u* is normalized w.r.t. the radius of the corresponding parallel, while the derivative with respect to *v* is normalized by a factor 1*/*2 as a meridian only covers half a circle. The normalized derivatives are represented in Supplementary Fig. 8(d). The impact of the regularization term on surface is discussed in Supplementary Section .

### Inference pipeline

#### Global approach with the CNN

The set *{****x***_*q*_*}*_1*≤q≤Q*_ of co-ordinates of the local maxima of the predicted score map 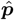 having a score greater than a chosen threshold *γ >* 0 are extracted, to obtain a single location per object in the input image ***I***. These voxels are assumed to represent the barycenters (or the voxels closest to them) of the objects of interest. A surface parameterized by 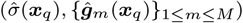, *{****ĝ***_*m*_(***x***_*q*_)*}*_1*≤m≤M*_ ) is generated for each estimated location ***x***_*q*_ .

In Nagini3D, the extraction of the local maxima, is processed by applying convolutions using appropriate kernels^6^ to perform the comparison between neighboring voxels (see Supplementary ). The efficient handling of convolution by GPUs makes this step very fast, explaining the noticeable gain in inference time proposed by **Nagini3D** compared to competing methods (last row of Table 1).

As the localization process can generate errors, a Non-Maximum-Suppression (NMS) is applied, which evaluates the Intersection-over-Union (IoU) between each pair of surfaces and removes potential duplicates representing the same object. Segmentation masks are generated by filling the interior of each surface. **Nagini3D** inference pipeline is summarized in Fig. 1.

#### Local approach (refinement step)

To enhance the result predicted by the network, an additional but optional step is added, that optimizes the position of each surface individually by iterativaly minimizing an energy functional based on image gradient through a gradient descent. To compute the energy a function, denoted *V*, convolves the image with a Gaussian kernel and computes the inverse of the gradient magnitude^7^. The resulting potential *V* (***I***) has a low intensity on the contour of the objects (where the norm of the gradients of the initial image are high) and high intensity anywhere else.

The energy for a surface ***ŝ*** parameterized by 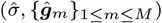 is defined by:

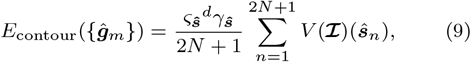

where *V* (***I***)(***x***) corresponds to the evaluation of potential *V* (***I***) at voxel ***x***. The energy corresponds to the evaluation of *V* (***I***) on the sampling of surface ***ŝ***. The first multiplicative factor *ς*_***ŝ***_ weights the energy depending on image intensity as follows:

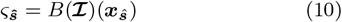

where *B*(***I***) is a convolved version of ***I*** with a smoothing gaussian kernel and ***x***_***ŝ***_ is the localized position of ***ŝ***. This factor reduces the impact of refinement for low contrasted objects for which boundaries are likely to be poorly captured by image gradient. For dataset in which all objects have similar intensity (CAPS, BONES), we cancel its impact by setting *d* = 0, for datasets with a wide range of object intensity (CREST), we set *d* = 3.

The second factor *γ*_***ŝ***_ weights the energy depending on object density in ***ŝ*** vicinity:

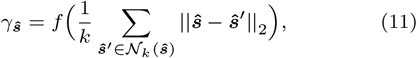

where *N*_*k*_(***ŝ***) denotes the set of *k* Nearest Neighboring objects of ***ŝ*** and *f* : ℝ ↦ ℝ is an increasing function. We set *k* = 4 and *f* : *t* ↦ 1*/*(1 + *e*^*a*(*b−t*^)), a shifted sigmoid (with *a* = 0.15 and *b* = 30 in our experiments).

This multiplicative factors is motivated by the high efficiency of snake-based methods in optimizing surfaces in sparse regions, contrasted with their tendency to degenerate when applied to dense regions. In densely populated areas, where object boundaries are less distinct, optimization often leads to merging objects or collapsing certain structures. This adaptive parameter addresses the issue by modulating the optimization strength based on sparsity: the sparser the region, the more significant the refinement effect.

### Nagini3D parameters setting

The U-Net network considered has a number of layers of 3, each layer processes 2 convolutions, the activation functions are ReLU, it uses batch-normalization layers, the number of channels starts at 32 and is multiplied by 2 at each pooling step.

Using the regularization term, Nagini3D can be used with a high number of control points. For the experiments described in this paper the settings are: *K* = 8, *L* = 4 (i.e., *M* = 30), 2*N* + 1 = 101 (*N* = 50).

The training is performed with PyTorch, using AdamW optimizer with a learning rate of 5 *×* 10^*−*4^ (the other parameters are the default ones). The model is trained for 4000 epochs, the batch-size is 1, the patch-size of an image is 80 *×* 80 *×* 80. The patch are randomly flipped along each dimension for data-augmentation.

For the inference, the score and the NMS thresholds are selected automatically to maximize the accuracy over the validation set.

### State-of-the-art competing algorithms

Competing methods are divided into two categories:

- **Fully 3D methods** (including Nagini3D) analyze the entire volume. The related competitive methods are StarDist3D [31] (**SD3D**) and Omnipose [32] (**OP3D**), both trained on the respective datasets for each benchmark. These methods were trained on the dataset considered for the experiment following the author recommendations for training. A grid search hyper-parameters search was conducted to obtain the best model.
- **2.5D methods** analyze separately the 3D volume slice-by-slice and subsequently aggregate the results to reconstruct volumetric segmentation. The advantage of these methods is that they are trained on massive 2D annotated sets, which make them competitive requiring no additional training. The competing methods include Cellpose2.5D [24] (**CP2.5D**); CellposeSAM [28] (**CP-SAM**) based on Cellpose 2.5D but using pre-trained transformer for enhanced performance; CellStitch [30], combined with both Cellpose2D [24] (**StitchCP2D**) and StarDist2D [25] (**StitchSD2D**); and u-Segment [29] (**uSeg2.5D**), which natively employs Cellpose2D. These methods use the pre-trained models of StarDist2D and Cellpose2D/CellposeSAM that were trained on massive datasets to make them as versatile as possible. All the models available were tested, with multiple hyper-parameters settings for inference. The results displayed are the best result among all the tested configurations.

### Datasets for experiments

Images used to benchmark the algorithms come from the three following datasets:

- **BONES** is a dataset of 28 images of fluorescence microscopy depicting nuclei of bone cells cultivated on furrows of various angles to investigate the impact of substrate tilt on cellular development. The set is split into 18 volumes for training, 3 for validation and 7 for testing. Each 3D volume (200 *±* 120 *×* 180 *×* 310 voxels) (the first dimension depends on the furrow angle) contains approximately 240 objects. The GT masks were segmented automatically and curated by an expert. Courtesy of SAINBIOSE Laboratory (Mines Saint Etienne, University Jean Monnet Saint-Etienne, INSERM, U1059, France). Example of slice and volume are illustrated in Supplementary Fig. 8(a).
- **CAPS** is a synthetic dataset, depicting spherical caps (portion of sphere cut off by a plane). The geometry of these objects could be assimilated to those of erythrocytes (red blood cells). Caps of various sizes and thicknesses randomly cut off represent both star-convex and non star-convex objects. The proportion of non-star-convex objects is driven by the ranges of possible positions of the cut-off plane and of possible thicknesses. The thicker the sphere is and the smaller the part of the sphere the cut-off plane keeps, the more likely the corresponding cap is to be star-convex. Blur and Poisson-Gaussian noise were added to simulate the imaging conditions in fluorescence microscopy images. The set contains 10 training images, 3 for validation, 5 for testing. Each image (200 *×* 200 *×* 200 voxels) contains approximately 220 objects. Example of slice and volume are illustrated in Supplementary Fig. 8(a).
- **CREST** is a real set containing 10 training images, 5 for testing and 3 for validation representing zebrafish trunk neural crest cells (TNCs). The mCherry fluorescence of neural crest nuclei was from the transgenic zebrafish line Sox10:mG[48]. Each image (*≈*90 *×* 500 *×* 1000 voxels) contain approximately 200 objects. This dataset depicts TNCs, which move from wide progenitor regions into narrow (*≤*3 µm) channels between the surrounding tissues (notochord, neural tube and somites) during embryonic development at 22hpf. Courtesy of Scarpa Lab (Department of Physiology, Development and Neuroscience, University of Cambridge). Example of slice and volume are illustrated in Fig. 5 and Supplementary Fig. 8(a).

### Metrics

The tested methods were evaluated following the method presented in [31]: predicted objects are paired with the GT by solving an assignment problem maximizing the sum of the IoU of all matches. A GT cell with no corresponding prediction is considered a false-negative (*FN* ) and a predicted cell with no corresponding GT is considered a false-positive (*FP* ). A predicted cell having a corresponding GT is considered a truepositive (*T P* ) prediction and the corresponding IoU is computed. To each IoU threshold *τ ∈* [0, 1] is associated:

- *T P* (*τ* ), the number of TP with an IoU higher than *τ*,
- *FN* (*τ* ) the number of GTs with no matching prediction or in a pair with an IoU smaller than or equal to *τ*,
- *FP* (*τ* ) the number of predicted cells with no matching GT or in a pair with an IoU smaller than or equal to *τ* .

These notations are essential to define the following four metrics:

- Recall: 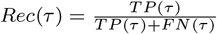 measures the proportion of correctly segmented objects (at a given IoU threshold *τ* ) among all objects to be detected;
- Precision: 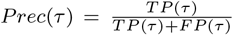 measures the proportion of correctly segmented objects (at a given IoU threshold *τ* ) among all objects predicted by the network;
- Accuracy: 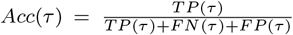 two previous metrics; aggregates the
- Score: 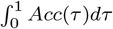 integrates the accuracy curve to get a summarizing metric that provides information on both detection and segmentation.

*Rec*(0), *Prec*(0) and *Acc*(0) are considered to evaluate the detection task alone (without considering the segmentation quality). Indeed, these values are computed considering any pairing with an IoU strictly higher than zero, meaning objects are localized at the right position regardless of the segmentation quality. Values of *Rec*(*τ* ), *Prec*(*τ* ) and *Acc*(*τ* ) with *τ >* 0 take the segmentation quality in consideration.

To assess the accuracy of segmentation boundaries relative to ground truth, we employ the Boundary Intersection over Union (BIoU) metric [49], which focuses specifically on boundary alignment. Unlike the standard IoU (which evaluates segmentation quality volumetrically and is thus insensitive to boundary errors in large objects) the BIoU restricts evaluation to boundary voxels only. Boundaries are extracted via binary erosion: the ground truth and predicted masks undergo three iterations of erosion (using a 1-voxel cube connectivity), and boundary voxels are defined as the difference between the original mask and its eroded version. This metric provides a more sensitive measure of boundary precision, particularly for large objects where boundary voxels constitute a small fraction of the total volume.

## Code availability

All source code and examples are made publicly available at the Nagini3D GitHub repository https://github.com/QuentinRapilly/NAGINI-3D/.

## Data availability

The synthetic dataset CAPS is available on the Zenodo website: https://zenodo.org/records/14931808. The BONES and CREST datasets were provided by courtesy of SAINBIOSE Laboratory (Mines Saint Etienne, University Jean Monnet Saint-Etienne, INSERM, U1059, France) and Scarpa Laboratory (Department of Physiology, Development and Neuroscience, University of Cambridge, UK), respectively.

## Acknowledgment

We would like to thank Leila Muresan (Cambridge Advanced Imaging Centre, UK) for the helpful computing resources (EP-SRC EP/Y008715/1) provided for the experiments. We are very grateful to Leila Muresan and Jérôme Boulanger (Laboratory of Molecular Biology, Cambridge, UK) for the fruitful discussions during the development of the method. We acknowledge Elena Scarpa (the Department of Physiology, Development and Neuroscience of the University of Cambridge, UK) for access to the dataset, named CREST in the manuscript, manually annotated by You-Hsuan Liu. You-Hsuan Liu is the recipient of a Cambridge Trust Taiwan-Cambridge PhD Scholarship. Pierre Maindron and Guénaëlle Bouet (from SAINBIOSE lab, Saint-Etienne, France) provided the dataset of real fluorescence microscopy images named BONES in this manuscript. Violette Thermes and Jérôme Bugeon (from INRAE, LPGP, Rennes, 35000, France) provided the zebrafish oocyte dataset used for the preliminary work depicted in the Supplementary information. We finally thank Sebastien Herbreteau (Ensai, CREST, Rennes, France) for the careful proof reading of the manuscript.

## Funding

This work was funded by Inria/SAIRPICO Team, a joint research team between Inria Center at University of Rennes (France) and Institut Curie, Chemical Biology of Cancer Unit, Paris (France).

## Author Contributions

Conception: Q.R.

Method implementation: Q.R.

Investigation and analysis: Q.R., A.B. and C.K.

Napari Graphical user interface (GUI) development: Q.R.

Data generation: Q.R. (CAPS set), Y.H.L. (CREST set).

Supervision: A.B. and C.K.

Funding acquisition: A.B. and C.K.

Writing-original draft: Q.R. and C.K.

Writing-review and editing: all authors.

## Competing Interest

The authors declare no competing interests.

## SUPPLEMENTARY INFORMATION

### Supplementary Methods

#### Exponential B-spline

The exponential B-Spline used for the surface parametrization is described as follows:

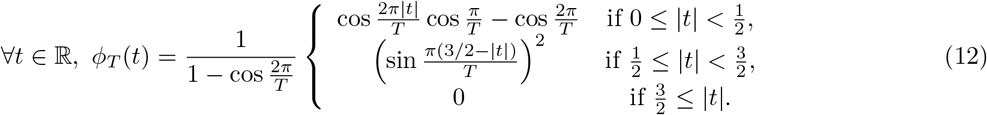

This spline perfectly reproduces ellipsoids for *T ≥* 3.

#### Closure and smoothness constraints

The constraints to guarantee that a surface described by Eq. 1 are closed and smooth (*C*^1^) are as follows:

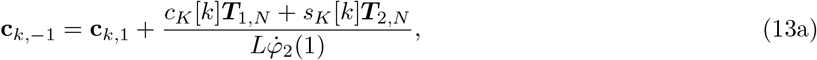

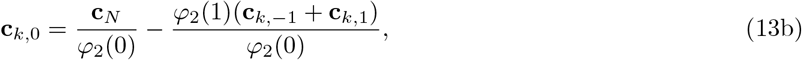

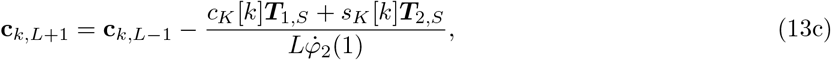

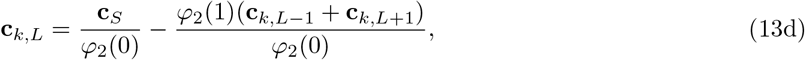

#### From generic to closed and smooth parametric surface

We show how to transition from the set of parameters 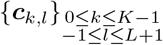 describing generic surfaces to the set *{****g***_*m*_*}* _1*≤m≤M*_ restraining the generated surfaces to closed and smooth ones. We rewrite the general equation of surface (the scale factor *σ* is omitted as it does not influence the demonstration):

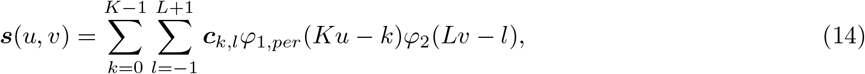

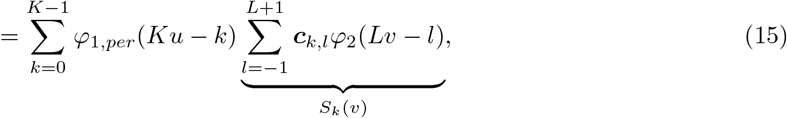

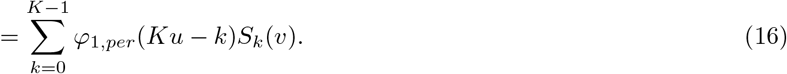

We then explicit the *S*_*k*_(*v*) term to make the *{****g***_*m*_*}*_1*≤m≤M*_ appear:

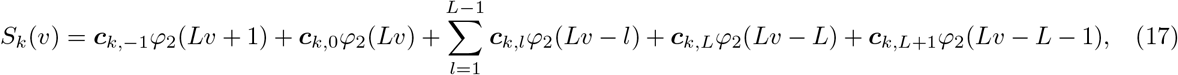

With

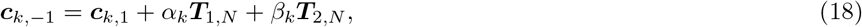

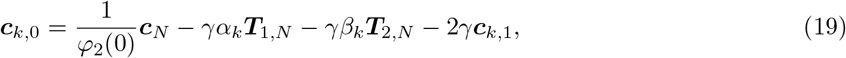

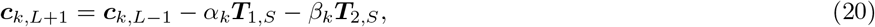

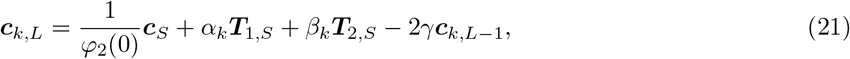

with 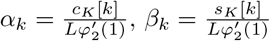, and 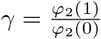. We obtain:

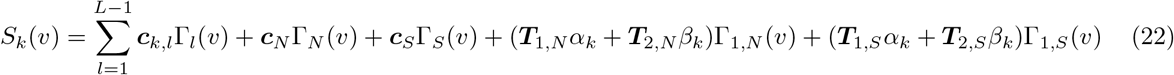

with

- Γ_1_(*v*) = *φ*_2_(*Lv −* 1) + *φ*_2_(*Lv* + 1) *−* 2*γφ*_2_(*Lv*),
- Γ_*L−*1_(*v*) = *φ*_2_(*Lv − L* + 1) + *φ*_2_(*Lv − L −* 1) *−* 2*γφ*_2_(*Lv − L*),
- 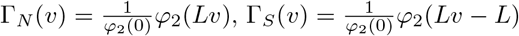,
- Γ_1*N*_ (*v*) = *φ*_2_(*Lv* + 1) *− γφ*_2_(*Lv*),
- Γ_1*S*_(*v*) = *γφ*_2_(*Lv − L*) *− γφ*_2_(*Lv − L −* 1),
- *∀l ∈ {*2, …, *L −* 2*}*, Γ_*l*_(*v*) = *φ*_2_(*Lv − l*).

We can finally rewrite the surface equation with the *{****g***_*i*_*}*_1*≤i≤I*_ by injecting (22) in (16):

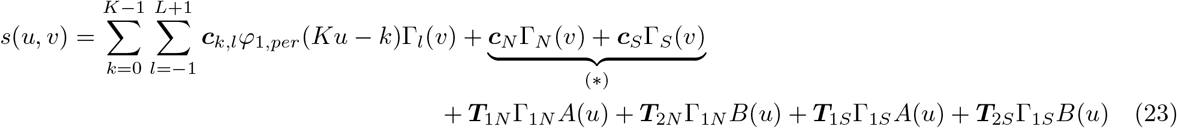

with 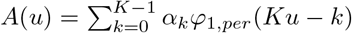 and 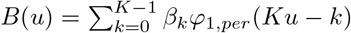. In (*\**) there is no term depending of *u* because with partition of unity properties of the basis functions this part is equal to 1. We can finally write:

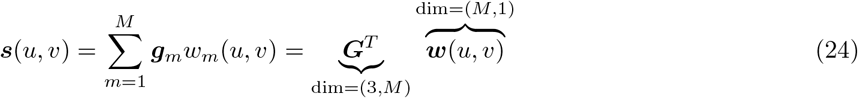

where

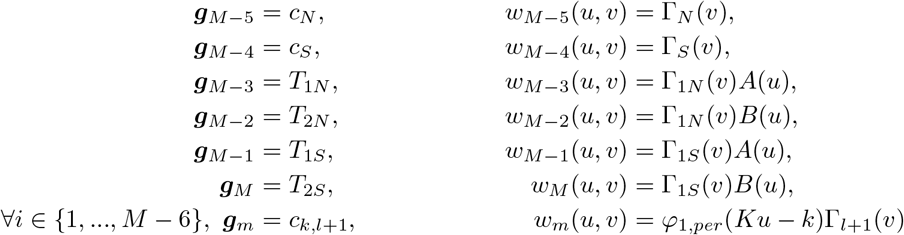

with *k* the quotient and *l* the remainder in the euclidean division of (*m −* 1) by (*L −* 1).

#### Fibonacci Lattice

A Fibonacci lattice containing 2*N* + 1 points 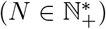, gives the latitudes and longitudes 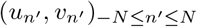 to use to sample the surface evenly as follows:

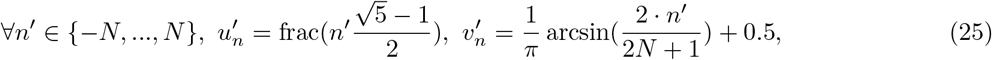

where frac : *x → x − ⌊x⌋* denotes the fractional part function. The sampling *Ŝ* = *{*ŝ_*n*_*}*_1*≤n≤*2*N*+1_, with 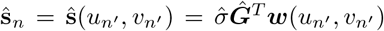 (where *n*^*′*^ = *n −N−* 1) samples the surface evenly as shown on 6b (c). The conventional sampling strategy regularly dividing *u* and *v* domains over-sample the poles as illustrated on Suppementary Figure 6a, because of the chosen surface parameterization.

**Figure 6.**
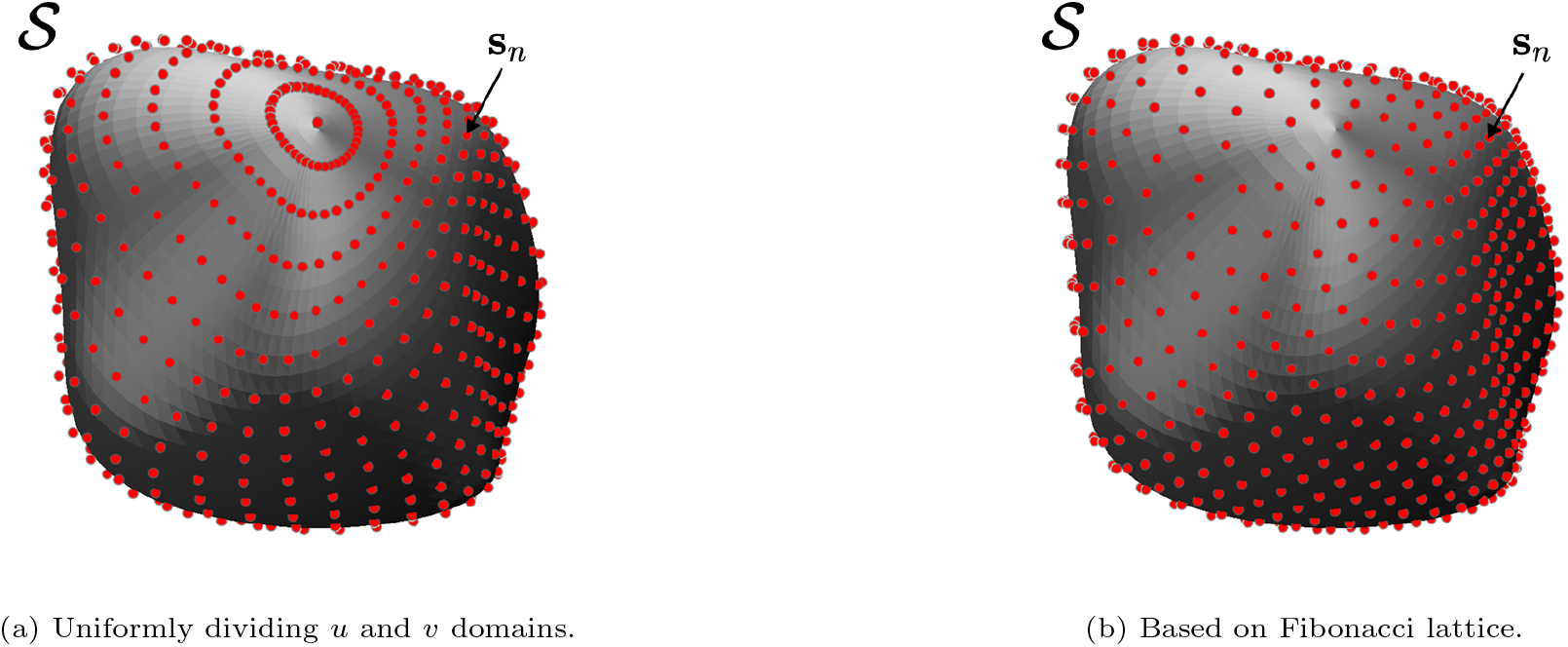
Impact of two different sampling strategies.

#### Geometry analysis

##### Local curvature

A measure of interest for further analysis of the object represented by surfaces is the mean local curvature. In some biological tasks, studying the curvature of objects can help to identify signs of specific biological events (e.g., cell division), to cluster objects according to their shape characteristics or evaluate the force applied on an object membrane.

The mean local curvature of 3D surfaces represented by Eq. (3), using *C* ^1^ basis functions, is defined as follows. The surface derivatives are used to define two vectors generating the tangent plane to the surface:

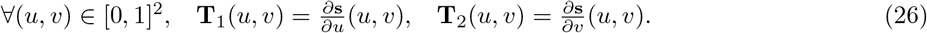

Then a vector normal to the surface can be defined by *∀* (*u, v*) ∈ [0, 1]^2^, **n**(*u, v*) = **T**_1_(*u, v*) ⊗**T**_2_(*u, v*) where ⊗ is the cross product in ℝ ^3^. Its normalized version is denoted 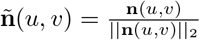.

The curvature is defined by 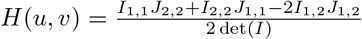 with:

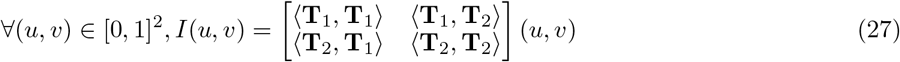

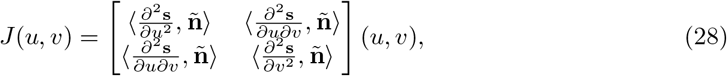

where *⟨·, ·⟩* denotes the Euclidean dot product on ℝ^3^.

In our case, we considered the mean local curvature dedicated to convexity concerns and of great interest to evaluate the force applied on object surfaces, but the Gaussian curvature can also be computed to identify saddle points. It is defined as:

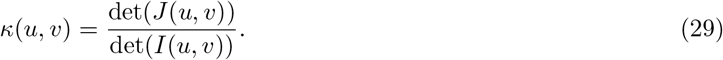

##### Principal Component Analysis

To perform the Principal Component Analysis (PCA) on the CREST dataset we followed the process illustrated in Figure 3: we simply transformed the parameter sets into 3*M* -dimensional vectors and extracted the mean shape along with the principal directions. It did not require alignment of the segmented objects. Indeed, when the localization step is executed correctly, objects are reconstructed at their centroid, eliminating the need for translation correction in space. Furthermore, although not theoretically guaranteed, we observe that the network consistently predicts object orientation: the north-south axis is generally aligned, and the remaining control points are arranged similarly around this axis in the absence of major inter-object deformation.

##### Inference details

The inference procedure implies to locate the object using the local maxima of the predicted score map 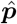. This step is done using convolutions with well chosen kernels comparing a voxel to its neighbors, to be efficiently computed on GPU. It makes our method the fastest among all the tested methods. The process is illustrated in Supplementary Figure 7.

**Figure 7.**
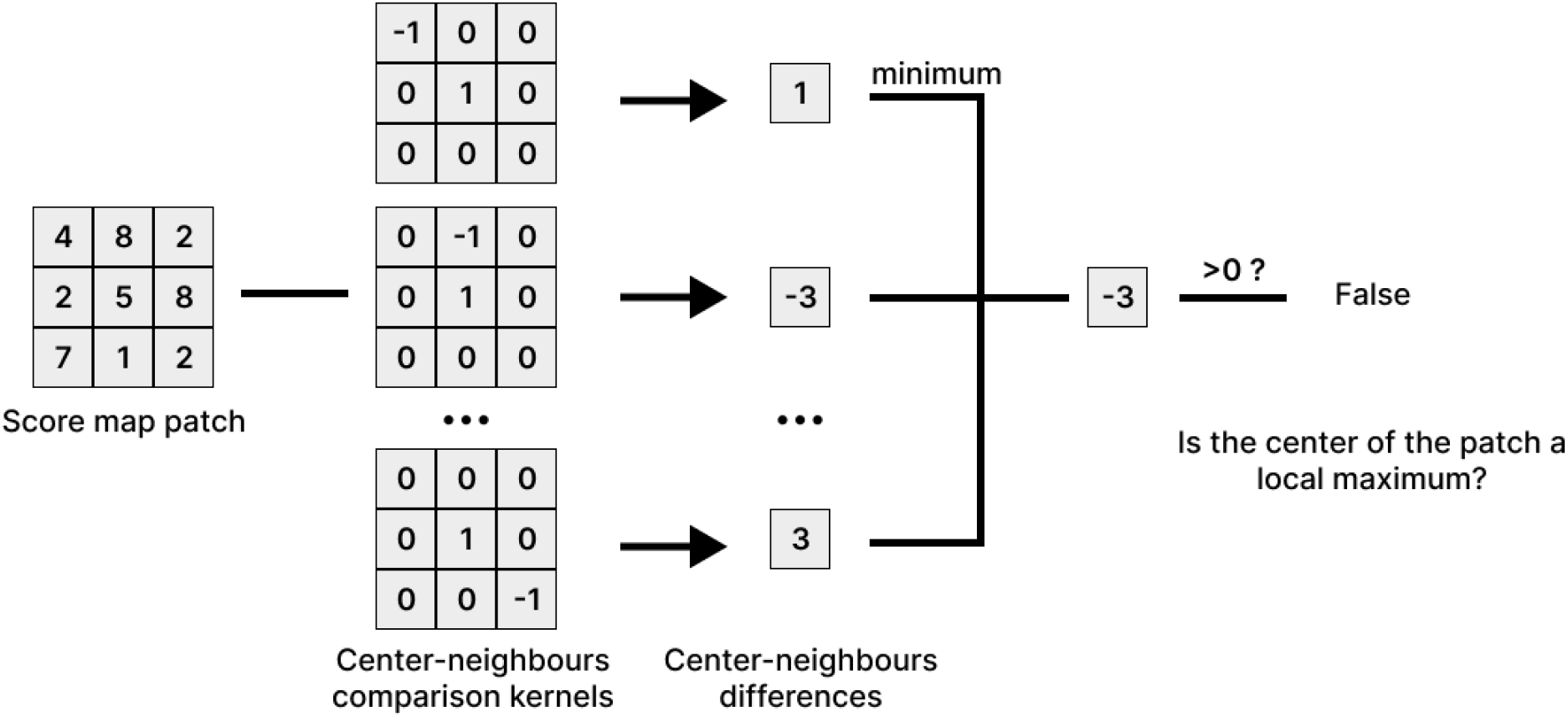
**Local maxima detection** using convolution kernels to compare each voxel to its neighbors. The process is illustrated in 2D but is equivalent in 3D.

### Supplementary Figures in Method Section

**Figure 8.**
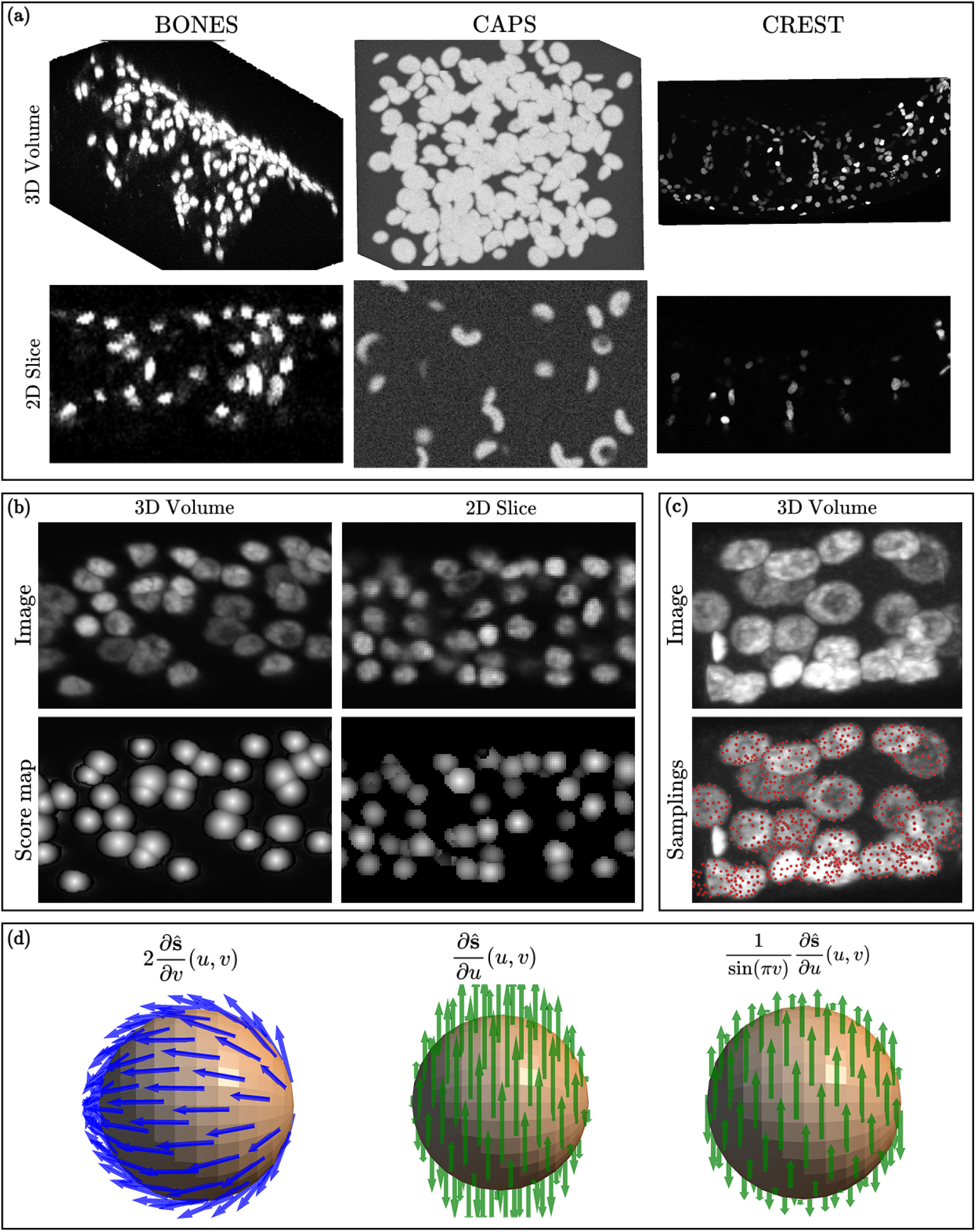
Ilustration of a few steps of Nagini3D (see workflow in Fig. 1). **(a)** Examples of volume and slice for each of the three datasets (BONES, CAPS, CREST), used to evaluate the segmentation methods. **(b)** Examples of volume and slice of the GT scores associated to input images. **(c)** Example of GT object samplings generated with the FPS algorithm. **(d)** Normalized and non-normalized surface derivatives computed on a spherical parametric surface.

#### Additional experiments

**Figure 9.**
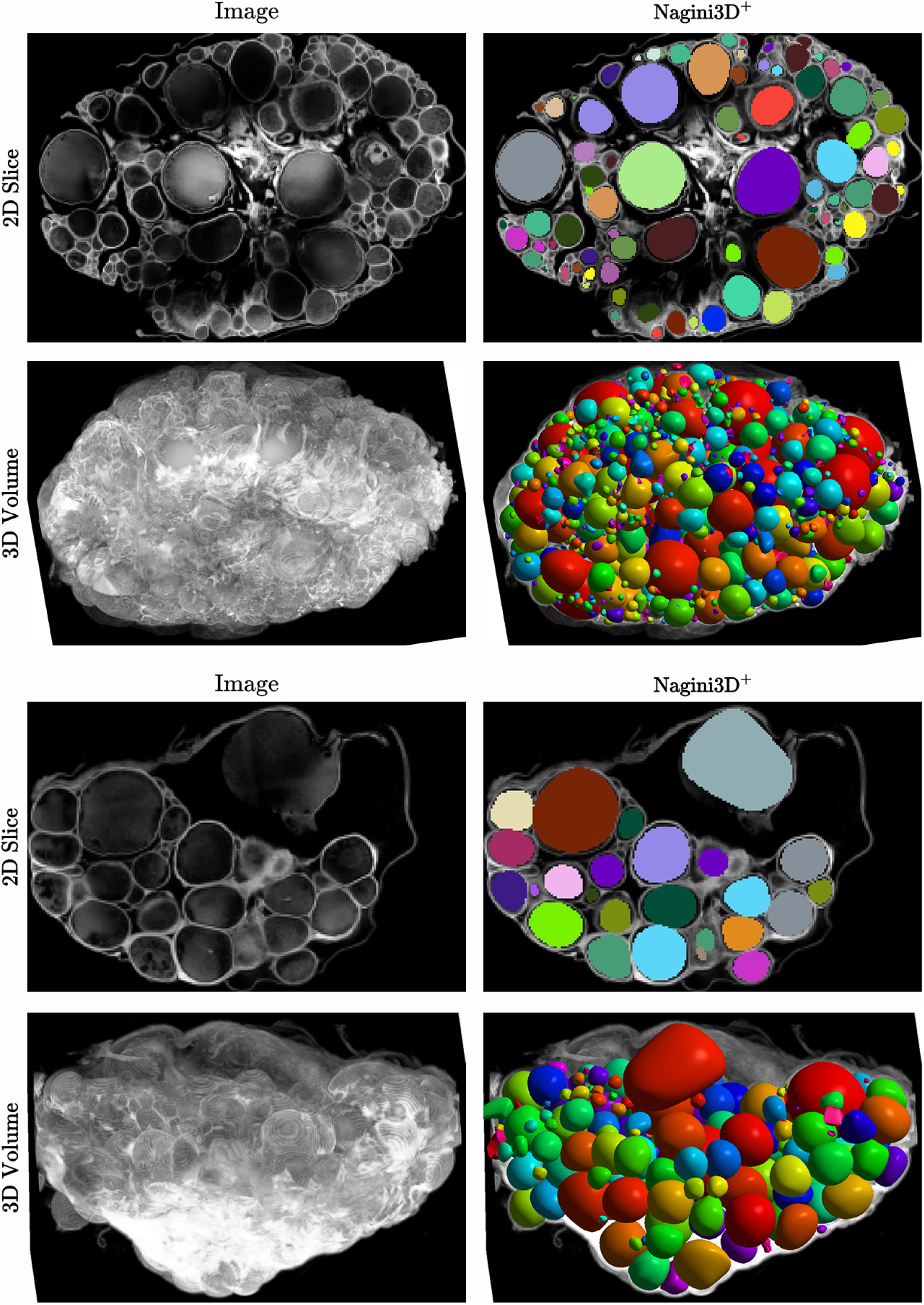
Segmentation conducted on zebrafish oocytes images. On challenging images using a membrane-specific staining and with touching object with very different scales, **Nagini3D**^+^ accurately segment most of the objects. Some of the smallest are missing.

##### Preliminary work on multiple scale object

Preliminary work was conducted on fluorescence images of zebrafish oocytes acquired with a confocal microscope. This dataset comprises 10 training images, 3 for validation, and 6 for testing. Volume sizes range from 150*×*150 *×* 150 to 250 *×* 500 *×* 320 voxels. Each image contains hundreds of roughly spherical objects with radii ranging from 4 to 20 voxels. These images present several notable challenges: the staining is membrane-specific, objects are densely packed and thus in contact, and the image contains objects of highly variable size (the volume of large objects can be up to 100 times greater than that of small ones). In this ongoing work, the goal is to demonstrate **Nagini3D**^+^ ability to process images with highly variable object sizes. In Supplementary Figure 9, we observe that despite cellular labeling, the majority of objects are segmented very accurately, except for a few of the smallest ones.

##### Nagini3D is suited for anisotropic dataset analysis

While 3D microscopic images are sometimes isotropic (where a voxel displacement corresponds to the same spatial shift along each dimension), it is common, depending on the microscope technology or its settings, for one dimension to represent a different spatial displacement than the other two. For such anisotropic images, an adaptation of the algorithm is required to account for the spatial discrepancies among the three axes.

The datasets used for the experiments in this paper are anisotropic or nearly so (when this is not the case, the anisotropy is mild and does not affect algorithm performance). Thus, the standard version of the algorithm was used. However, a version adapted for highly anisotropic data is available. The modifications are:

- flips along the three axes for data augmentation are no longer applied;
- the user specifies the anisotropy ratio for each axis;
- during pre-processing, when generating samples, the anisotropy ratio is used in “Farthest Point Sampling” to generate a uniformly distributed sampling as if the object were dilated to an isotropic space;
- the network is trained to predict surfaces that represent objects isotropically.

During inference, users can either reconstruct objects in the image space (with predicted objects simply rescaled along each axis) or perform segmentation in an isotropic space.

The FPS-specific technical step simply involves computing the most distant voxels based on geodesic distance, accounting for the fact that displacement is not equivalent across directions. Incorporating the anisotropy ratios for each dimension ensures that distances remain consistent with physical reality.

**Figure 10.**
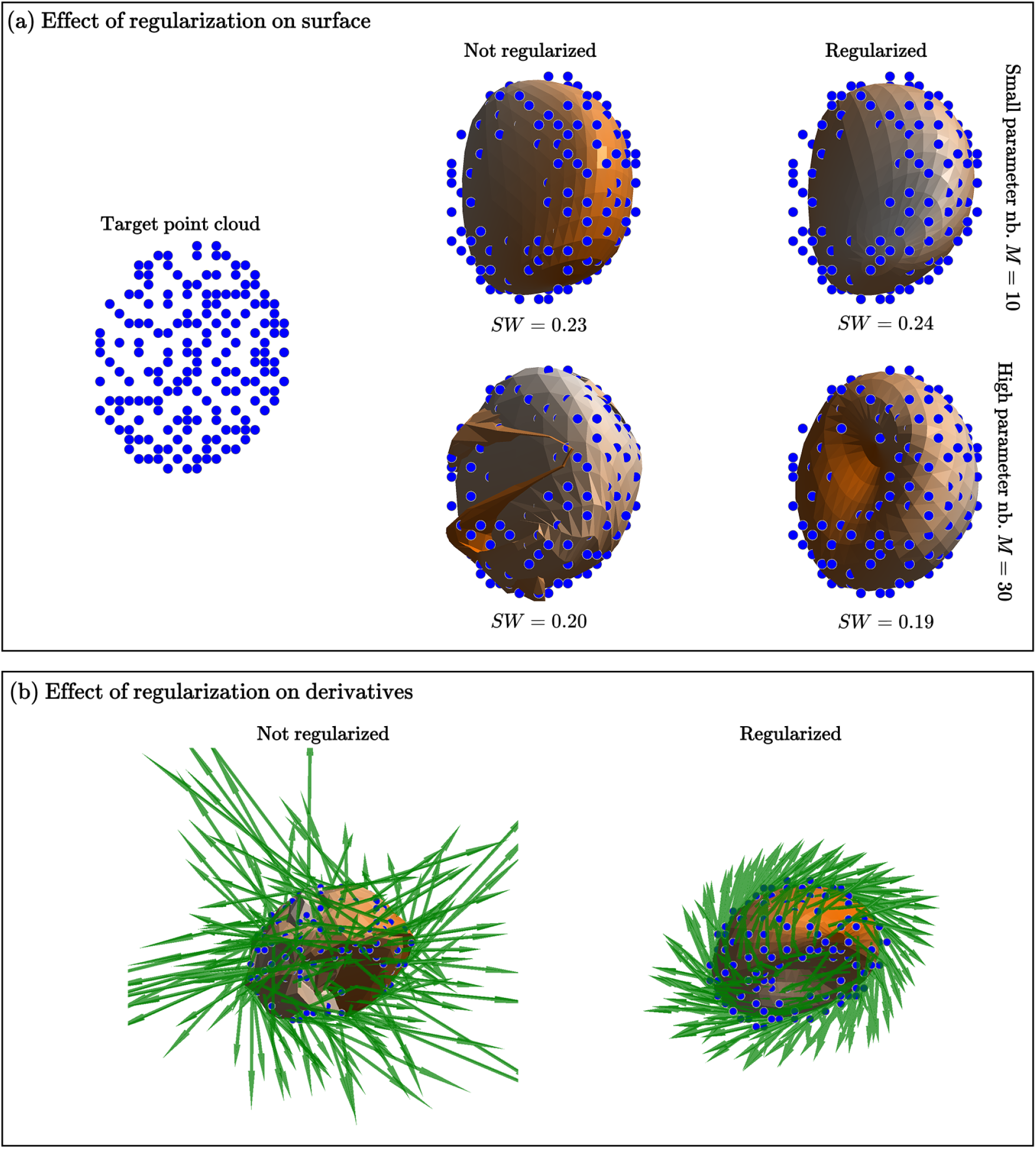
Impact of the regularization on target-surface matching. **(a)** Surface are optimized to match a target point cloud by minimizing Equation (5). Multiple settings are evaluated: with or without regularization; with a few (*M* = 10) or high (*M* = 30) number of surface parameters. The resulting data-attachment (Sliced-Wasserstein distance *SW* ) is shown, lower is better. **(b)** Impact of the regularization on the surface derivatives for the estimated surfaces with *M* = 30.

#### Surface regularization motivation

When designing this method, a first version without regularization was developed. The model can be trained to predict surfaces parameterized by a small number of control points. However, when the number of parameters is slightly increased, the network tends to predict degenerate, self-intersecting surfaces and never recover plausible surfaces throughout training. This is not a surprise as the random initialization of the network weights: consequently, surfaces predicted in early epochs are also random. A randomly initialized surface with many parameters is highly likely to be degenerate, whereas a surface with few parameters remains close to a plausible shape. To predict surfaces with a larger number of parameters while ensuring the plausibility of the resulting shapes, it is required to penalize self-intersecting shapes. Such shapes are typically non-smooth, exhibiting significant local stretching. Penalizing surface derivatives is therefore an effective solution.

The regularization term defined in Eq. (8) penalizes the norm of the surface derivatives and enables to train the network to predict surfaces parameterized with a high number of control points (reaching up to *M* = 46 (*K* = 10, *L* = 5) during our experiments). A display of the effect of the regularization term is illustrated in Figure 10(a). Surface are optimized to match a target sampling using Eq. (5) either with or without the regularization term. For a few parameters only (*M* = 10) the regularization seems to be of no interest as the surface optimized without is realistic as well. The impact arises for more parameters (*M* = 30) for which the surface trained without is completely unrealistic (self-intersecting), while the surface trained with regularization remains plausible.

As depicted in Supplementary Figure 10(b), the regularization term tends to smooth the surface, preventing stretched areas with high surface derivatives.

**Figure 11.**
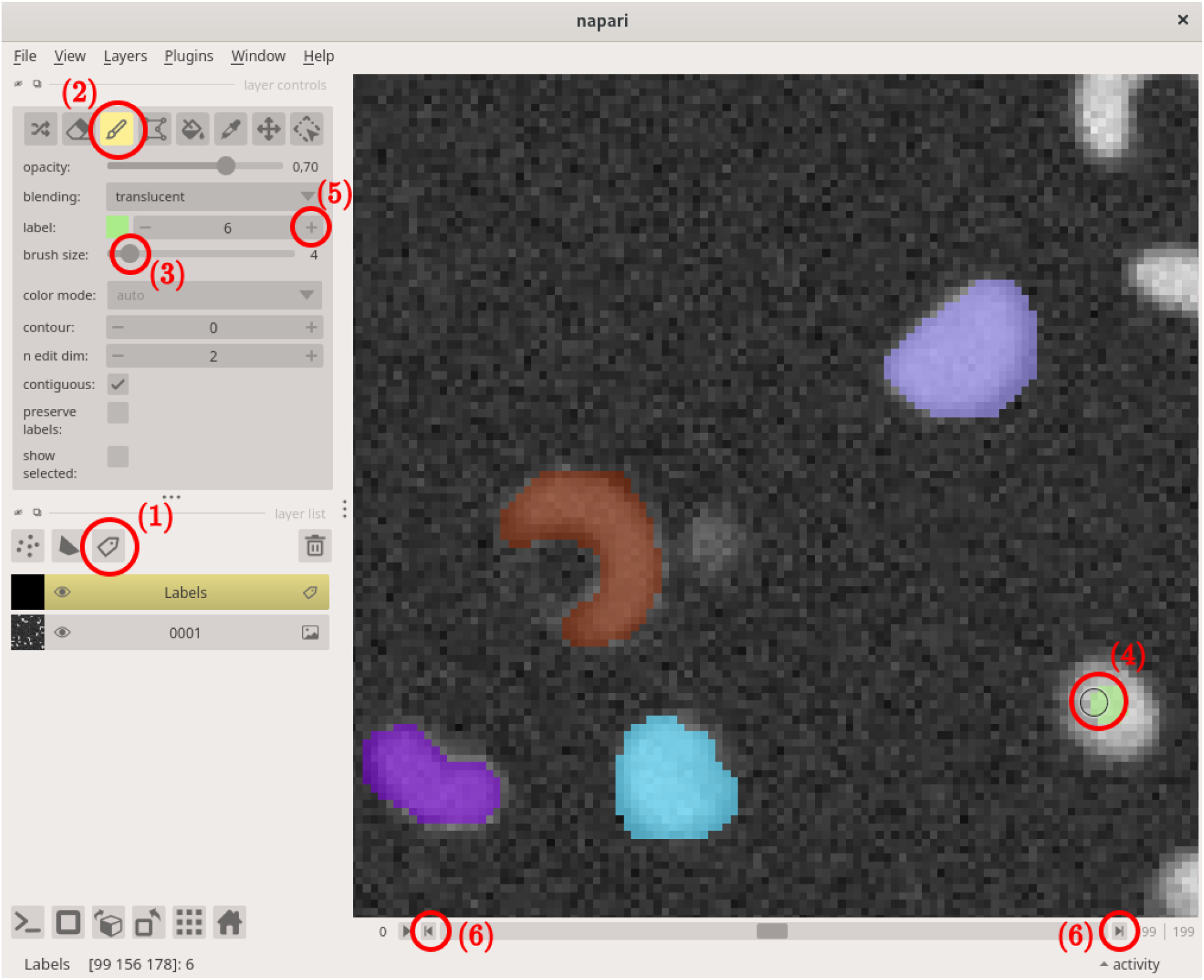
**Labeling with Napari software:** after drag-n-dropping your image into Napari, add a Labels layer **(1)**. Select the brush tool **(2)**. You can modify the brush size **(3)**. Annotate an object with your mouse cursor **(4)**. Once an object is completely annotated, switch to the following label **(5)**. Once on slice is completely annotated navigate to the next ones **(6)**. Take care to keep the same label from a slice to the following one for the same object.

#### Tutorials

##### Data annotation

The training pipeline requires to provide a mask labeling exhaustively each object of interest inside a corresponding volume. In what follows, we describe several methods to generate such masks.

To guarantee optimal performance, the masks used for training have to be annotated exhaustively: every object should be labeled. But, as mentioned in Section “Nagini3D only needs a few annotated samples”, the annotation can be coarsely performed as **Nagini3D** refinement will reduce network imprecision. However, the more accurate the GT masks, the better the segmentation quality.

###### Manual annotation

This method is the most time-consuming and tedious but also the most precise because the masks correspond exactly to what the end-user wish to extract.

Several software provide annotation tools like Napari, LabKit, QuPath, … To our opinion, the most convenient tool is Napari. We illustrate how to annotate a volume using Napari in Supplementary Figure 11.

###### Semi-automatized annotation

In order to save time but still have reliable ground-truth mask, the best strategy is to generate the segmentation masks automatically and manually curate them if required (e.g., with Napari tool) in order to eliminate potential segmentation errors generated by the automated method.

A good and very simple example is to use an Otsu thresholding available in Scikit-learn (see https://scikit-image.org/docs/0.25.x/auto_examples/segmentation/plot_thresholding.html) which often already provides satisfying results for segmenting bright objects against a dark background. This method provides a binary segmentation and must be post-processed to separate instance, for example with a watershed algorithm (see https://scikit-image.org/docs/0.25.x/auto_examples/segmentation/plot_watershed.html).

More sophisticated tools can be used to generate masks. For instance, the end-user can supply masks produced by efficient 2.5D method (e.g., Cellpose-SAM) to **Nagini3D**.

## Footnotes

1 “Classical” or “old-school” methods refer to methods that are not based on deep-learning.

2 More details on CAPS and CREST datasets in the “Data” paragraph of Method Section

3 In the **Nagini3D** training procedure, surface estimation is unaffected, but object localization degrades if labels are missing.

4 Stored using *numpy*.*uint16* type to minimize the weight.

5 The amount of GPU memory used depends on object density as a part of the loss is computed only on foreground voxels.

6 A kernel computes the difference between the center value and one of the neighbors.

7 *V* corresponds to the function *inverse gaussian gradient* from **Scikit-image** python package [47]

## Notes

https://github.com/QuentinRapilly/NAGINI-3D

### Competing Interest Statement

The authors have declared no competing interest.

